# Effects of early life adversity on maternal effort and glucocorticoids in wild olive baboons

**DOI:** 10.1101/2020.12.11.418095

**Authors:** Sam K. Patterson, Katie Hinde, Angela B. Bond, Benjamin C. Trumble, Shirley C. Strum, Joan B. Silk

## Abstract

Adverse experiences during early life exert important effects on development, health, reproduction, and social bonds, with consequences often persisting across generations. A mother’s early life experiences can impact her offspring’s development through a number of pathways, such as maternal care, physiological signaling through glucocorticoids, or even intergenerational effects like epigenetic inheritance. Early life adversity in female yellow baboons (*Papio cynocephalus*) predicts elevated glucocorticoids, reduced sociality, shortened lifespan, and higher offspring mortality. If baboon mothers with more early life adversity, experience poorer condition and struggle to provide for their offspring, this could contribute to the persisting transgenerational effects of adversity. Here, we examined the effects of mothers’ early life adversity on their maternal effort, physiology, and offspring survivability in a population of olive baboons, *Papio anubis.* Mothers who experienced more adversity in their own early development exerted greater maternal effort (i.e., spent more time nursing and carrying) and had higher glucocorticoid metabolites than mothers with less early life adversity. Offspring of mothers with more early life adversity had reduced survivability compared to offspring of mothers with less early life adversity. There was no evidence that high maternal social rank buffered against the effects of early life adversity. Our data suggest early life experiences can have lasting consequences on maternal effort and physiology, which may function as proximate mechanisms for intergenerational effects of maternal experience.

## Introduction

Early life environments can have profound and lasting consequences. In humans, exposure to early adversity increases susceptibility to a variety of health problems, including cardiovascular disease, diabetes, obesity, and renal failure in adulthood (Barker, Eriksson, Forsen, & Osmond, 2002; Gluckman, Hanson, Cooper, & Thornburg, 2008). Studies of nonhuman animals have found an effect of early life adversity on adult physiology, sociality, fecundity, and survival (Descamps, Boutin, Berteaux, McAdam, & Gaillard, 2008; Douhard et al., 2014; Lea, Altmann, Alberts, & Tung, 2015; Monaghan, 2008; Nussey, Kruuk, Morris, & Clutton-Brock, 2007; Petrullo, Mandalaywala, Parker, Maestripieri, & Higham, 2016; Pigeon & Pelletier, 2018; Tung, Archie, Altmann, & Alberts, 2016). Adverse experiences, including harsh ecological and social conditions, in early development can exert long-term effects that transfer to the next generation, although the nature of the effects varies across taxa. For example, mothers’ early life adversity may be associated with greater offspring size and faster offspring growth (chicken: Lindqvist et al., 2007; Goerlich, Nätt, Elfwing, Macdonald, & Jensen, 2012; vole: Helle, Koskela, & Mappes, 2012; cichlid: Taborsky, 2006; fruit fly: Vijendravarma, Narasimha, & Kawecki, 2010); smaller size or slower growth (water flea: Andrewartha & Burggren, 2012; zebra finch: Naguib & Gil, 2005), and or have mixed effects (butterfly: Saastamoinen, Hirai, & van Nouhuys, 2013; hamster: William Huck, Labov, & Lisk, 1986, 1987; reviewed in Burton & Metcalfe, 2014). A mother’s own early life adversity has also been linked to offspring physiology, immunity, reproductive success, personality, and survival in short-lived captive animals (Burton & Metcalfe, 2014) as well as survivorship in several long-lived wild primates (Zipple et al., 2020; Zipple, Archie, Tung, Altmann, & Alberts, 2019).

A mother’s own early life experiences can affect her offspring through a number of pathways, as these experiences influence her adult phenotype, and thereby impact her offspring (Kuzawa, 2005; J. C. Wells, 2014; J. C. K. Wells, 2003, 2010). Maternal genes and epigenetic modifications can substantially shape offspring phenotype. Early life experiences can shape a mother’s biology and health by inducing epigenetic modifications (Conching & Thayer, 2019; Jablonka & Raz, 2009; Kuzawa & Thayer, 2011). For example, a low-protein maternal diet during gestation resulted in epigenetic silencing of a gene associated with type 2 diabetes risk in rats (Sandovici et al., 2011). A growing body of research demonstrates early life effects can be transferred to offspring via germline epigenetic inheritance (Conching & Thayer, 2019; Jablonka & Raz, 2009; Kuzawa & Thayer, 2011). For example, male mice exposed to early separation from their mothers experienced epigenetic changes in their sperm, and similar epigenetic changes were found in the neurons of the exposed males’ female offspring (Franklin et al., 2010).

In addition to germline epigenetic inheritance, maternal behavior and physiology can also have intergenerational impacts on offspring. Maternal care, hormonal signaling, and overall maternal condition shape a suite of offspring outcomes including health, cognitive development, dispersal patterns, and reproductive strategies (Bernardo, 1996; Catalani, Alemà, Cinque, Zuena, & Casolini, 2011; Groothuis & Schwabl, 2008; Langley-Evans, 2007; Mateo, 2014; Mousseau & Fox, 1998; Sheriff & Love, 2013). Meta analyses across 151 studies suggest that maternal effects account for half as much phenotypic variation as do additive genetics among short-lived vertebrate and invertebrates (Moore, Whiteman, & Martin, 2019). In birds, rodents, and primates, maternal presence and the type of maternal care behavior received as a neonate is linked to the type of maternal care that the individual provides to its own future offspring (Dettmer, Heckman, Pantano, Ronda, & Suomi, 2020; Doumas, Margolin, & John, 1994; Lynn A Fairbanks, 1996; Francis & Meaney, 1999; Müller et al., 2011; Sproul Bassett et al., 2020). How other dimensions of a mother’s early life experiences affect the maternal care and investment she provides is less clear. Early life adversity is linked to altered hypothalamic pituitary adrenal (HPA) function in adult monkeys and humans (Anacker, O’Donnell, & Meaney, 2014; Palma-Gudiel, Córdova-Palomera, Leza, & Fañanás, 2015; Petrullo et al., 2016; Rosenbaum et al., 2020; Tyrka, Ridout, & Parade, 2016), but more work is needed to determine how a mother’s own early life experiences affect the GC concentrations she transfers to her offspring through the placenta and through her milk. Women exposed to early life adversity have smaller bodies, ovaries, and uteruses when they begin to reproduce, and produce smaller offspring than women who are not exposed to early adversity (Ibáñez, Potau, Enriquez, & De Zegher, 2000; Martorell, Ramakrishnan, Schroeder, & Ruel, 2009; Ramakrishnan, Martorell, Schroeder, & Flores, 1999; Stein, Zybert, van de Bor, & Lumey, 2004). In yellow baboons, offspring born to mothers who themselves experienced early maternal loss have an elevated mortality risk and their deaths often precede their mothers’ death by a year or two, suggesting that these mothers struggle to meet the needs of their growing offspring (Zipple et al., 2019). If a mother’s own early life adversity constrains her ability to invest in her offspring and affects the behavioral or physiological signals she sends to her offspring, this could shape offspring phenotype and development.

Existing work demonstrates a connection between early life adversity experienced by the mother and her offspring’s outcomes. But it is not entirely clear if these intergenerational effects are the result of epigenetic transmission (Conching & Thayer, 2019; Jablonka & Raz, 2009; Kuzawa & Thayer, 2011), variation in maternal care and signaling as a function of the mother’s own early or later life adversity, direct effects of the current environment on offspring, or a combination of these mechanisms. Parenting behavior, as opposed to epigenetic transmission or in-utero investments, is the primary mechanism driving intergenerational effects of maternal presence in captive rhesus macaques (Dettmer et al., 2020). In humans, variation in early life adversity is often confounded with later life adversity such as access to healthcare, night-shift work, and diet (Snyder-Mackler et al., 2020). To disentangle these factors, a system free from these confounds is needed. Long-lived wild animals provide such a system and can thus serve as an important model species. In an effort to fill the gap in the existing literature, here we report the impact of early life adversity on maternal effort and physiology of wild multiparous olive baboons, *Papio anubis*. We also investigate the impact of maternal early life adversity on offspring survival in an attempt to replicate previous findings in primates (Zipple et al., 2020, 2019). Following previous work, we take advantage of long-term demographic and ecological data to assess several forms of early life adversity.

To assess maternal effort, we examine time spent nursing and carrying offspring, as well as maternal fecal glucocorticoid metabolite (GCM) levels. We use nursing and carrying as behavioral proxies for maternal effort because these are the most energetically demanding components of care for primate mothers (Jeanne Altmann & Samuels, 1992; Ross, 2001). Studies of the long-term consequences of nursing and carrying behavior on offspring are rare, but suckling behavior affects growth and survival in mountain goats (*Oreamnos americanus*) (Théoret-Gosselin, Hamel, & Côté, 2015) and it has been argued that infant-carrying has been conserved in primates because it reduces infant mortality risk (Ross, 2001). Maternal GCs reflect energy balance, stress, health, and fertility (Palme, 2019; Sapolsky, Romero, & Munck, 2000), allowing us to examine how early life adversity affects a mother’s ability to invest in offspring. Maternal GCs are transferred across the placenta and through mother’s milk (Meaney, Szyf, & Seckl, 2007; Pácha, 2000) and might also act as physiological signals and guide offspring development (J. C. Wells, 2014). Maternal-origin hormones are hypothesized to orchestrate offspring’s tradeoffs between developmental priorities in relation to maternal resources or environmental conditions (Allen-Blevins, Sela, & Hinde, 2015; K. Hinde et al., 2015). Elevated maternal GCs are associated with impaired offspring immune development, slower motor development, and less sociable temperament (reviewed in Lu et al., 2019). Offspring can use maternal GCs to guide their development in an adaptive way (e.g. B. Dantzer et al., 2013). For example, squirrel pups (*Tamiasciurus hudsonicus*) that experience high levels of maternal GCs, a signal of high population density, accelerate growth which improves their chance of survival (Dantzer et al., 2013). Rhesus macaque infants exposed to elevated maternal-origins GCs may prioritize somatic growth over behavioral development (K. Hinde et al., 2015).

We hypothesize that mothers’ own early life adversity will have a negative effect on their physiology and ability to invest in their offspring and this will negatively affect their offspring’s welfare. We test a number of predictions derived from this hypothesis:

1. Mothers who experienced more adversity during their own early development will produce offspring who nurse at higher rates. Early life adversity leads to poorer adult health and physical condition, and this is expected to predict reduced milk quality and quantity. Rhesus macaque mothers who experienced poor developmental conditions produce lower available milk energy (Pittet, Johnson, & Hinde, 2017). Reduced nutrient intake of lactating mothers results in lower milk yield (red deer: Loudon, McNeilly, & Milne, 1983; baboons: Roberts, Cole, & Coward, 1985; humans: (Brown, Akhtar, Robertson, & Ahmed, 1986; Emmett & Rogers, 1997), and lower milk yield is correlated with more suckling time (red deer: Loudon, McNeilly, & Milne, 1983; white-tailed deer: Therrien, Côté, Festa-Bianchet, & Ouellet, 2008).
2. Mothers who experienced more early life adversity themselves will carry offspring more. Although carrying offspring is energetically costly for mothers, transferring energy via milk to fuel the offspring’s independent locomotion is even more calorically demanding on the mother (Jeanne Altmann & Samuels, 1992). We therefore predict that mothers with more early life adversity will carry their offspring more than mothers with less early life adversity. Ventral carrying allows for suckling opportunities, aligning with Prediction 1.
3. Mothers who experienced more early life adversity will have higher GCM levels during pregnancy and lactation. Reduced nutrient intake and poorer energy balance are associated with higher GC levels (e.g., blue monkeys: Thompson, Higham, Heistermann, Vogel, & Cords, 2019, iguanas: Romero & Wikelski, 2001).
4. Mothers who experienced more early life adversity will have higher mortality among their offspring. In muriquis, blue monkeys, and yellow baboons, mothers’ early life adversity is associated with higher offspring mortality (Zipple et al., 2020, 2019).
5. High social status will buffer the effects of early life adversity. Female yellow baboons who experienced early life adversity showed greater reductions in fertility during drought years than females who were not exposed to early life adversity, but these consequences were eliminated if females were born to high ranking mothers (Lea, Altmann, Alberts, & Tung, 2015).

## Methods

### Study Site and Population

We studied four groups of wild baboons that range on the eastern Laikipia Plateau of central Kenya. These groups are monitored by the Uaso Ngiro Baboon Project (UNBP), directed by Dr. Shirley Strum. The study groups range in an area that is topographically diverse and averages 1718m above sea level. The habitat is dry savanna with grassy plains, acacia woodlands, and woodlands on the edge of dry sandy rivers. Annual rainfall is typically concentrated in two wet seasons (March-June, November-December; (Barton, 1993), though droughts are increasingly common). *Opuntia stricta*, an invasive non-indigenous cactus, has become an important part of the diet for all of the groups monitored by the UNBP (Strum, Stirling, & Mutunga, 2015). Access to the *Opuntia stricta* fruit has reduced seasonal variability in food availability and shortened interbirth intervals (Strum, unpublished data). Three of the study groups PHG, ENK, and YNT occupied overlapping home ranges and the fourth study group, NMU, ranged in a different area. Individuals in PHG, ENK, and YNT had more *Opuntia stricta* in their diet than those in NMU. From 2013-2017, the interbirth intervals for each study group are as follows: PHG 506±109.63 days (mean±sd), ENK 449.39±62.68 days, YNT 533±61.33 days, and NMU 566±87 days (*F*(3,67) = 8.065, p < 0.001; post-hoc tests show only a substantial difference between NMU and ENK: p < 0.001).

The troops we studied were descendants of two troops (PHG, MLK (formerly known as WBY)) that were translocated from the Rift Valley near Gilgil, Kenya to the Laikipia region in 1984 (Strum, 2005). PHG fissioned in a process that lasted from 2009 to 2011. The larger of the two daughter troops retained the name PHG and the smaller group was named ENK. PHG fissioned again in a process that lasted from 2010 to 2013.

Again, the larger of the two fission products retained the name PHG and was monitored through the end of the study period. The smaller group was named OGs and is not included in this study. In 2016, several females followed a natal male from PHG to ENK, and then left ENK to form a new group, YNT. The fourth troop we studied, NMU, is the product of a series of fusions between descendants of MLK and several indigenous troops.

Demographic records span the entire study period (Figure 1). Observers update demographic records daily and record when individuals are born, die, or disappear. Maternal kinship relationships among natal females were known from genealogical records extending back to the early 1970s. Data on herbaceous biomass are collected each month using the slanting pin intercept technique angled 65 degrees from vertical (McNaughton, 1979) and converted into biomass in gr/m2 using the adjusted equation HB=total hits X 0.847 (McNaughton, 1979; Western & Lindsay, 1984).

**Figure 1.**
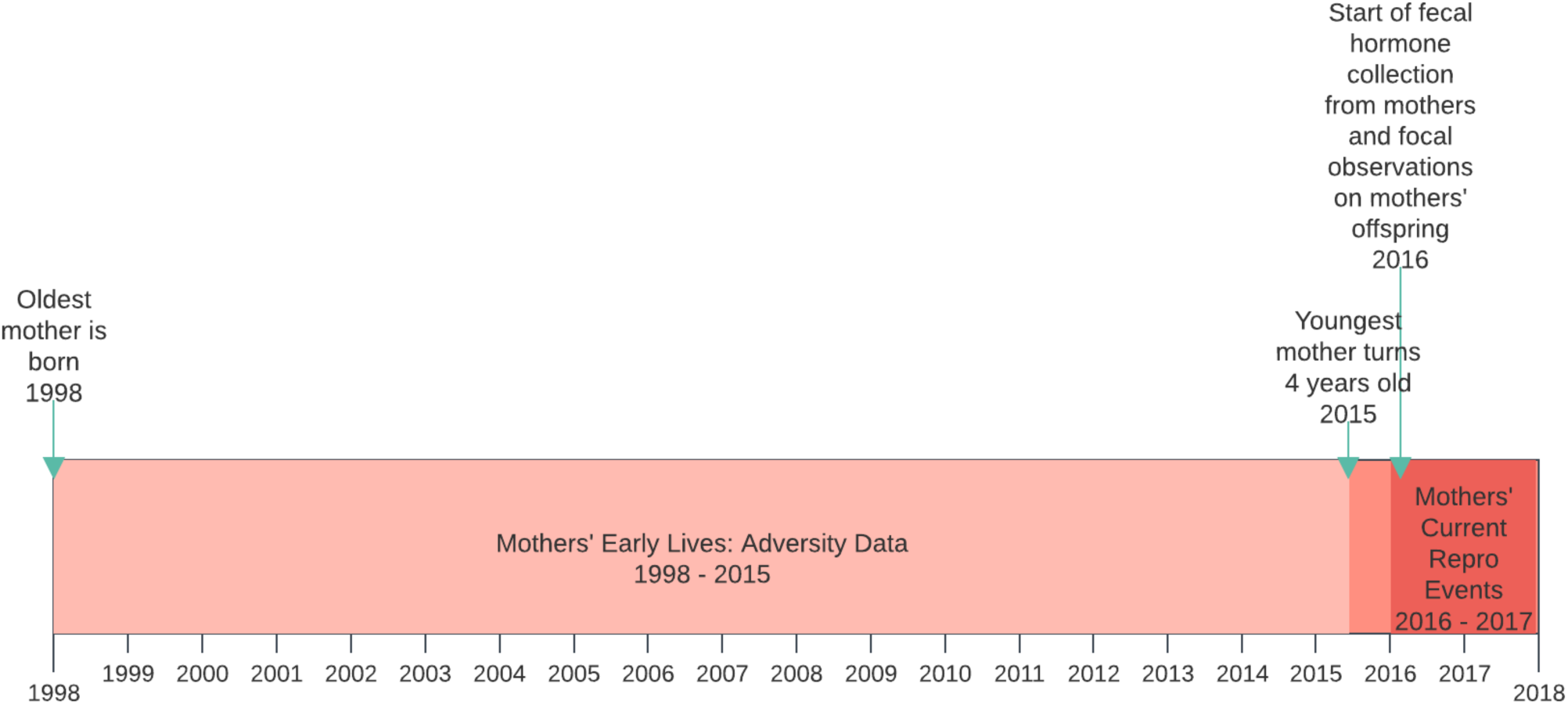
Study timeline

**Table 1.** Group composition at the beginning of the study period

|  | Adult/Subadult<br>Females | Adult/Subadult<br>Males | Juveniles | Infants | Total |
| --- | --- | --- | --- | --- | --- |
| PHG | 16 | 9 | 10 | 14 | 49 |
| ENK | 9 | 4 | 12 | 15 | 40 |
| YNT | 6 | 4 | 7 | 5 | 22 |
| NMU | 34 | 23 | 41 | 22 | 120 |

### Subjects

We conducted behavioral observations on 44 mothers and 47 infants from October 2016 to December 2017 (Figure 1). This sample represents all mother-infant dyads with infants under 1 year of age during the 2016-2017 study period. Dates of birth, rank, maternal kinship, and all components of early life adversity were known for 38 mothers. This study is restricted to multiparous mothers as the myriad complexities of primiparity (e.g., Carrera, Sen, Heistermann, Lu, & Beehner, 2020; Dettmer, Rosenberg, Suomi, Meyer, & Novak, 2015; K. J. Hinde, 2009; K. Hinde et al., 2015; Mas-Rivera & Bercovitch, 2008; Nuñez, Grote, Wechsler, Allen-Blevins, & Hinde, 2015; Pittet, Johnson, & Hinde, 2017) and small subset (N=7 out of 38 mothers) risked obscuring the early life adversity phenomena of immediate interest. The final behavioral dataset included 31 mothers and 34 offspring. Offspring mortality outcomes were available for all multiparous mothers in our sample. The mortality dataset includes 80 offspring, of which 10 died during infancy.

### Behavioral Observations

Observers conducted approximately 3900 complete 15-min focal samples during the 15-month study period on all infants under one year of age. Each of the 34 focal offspring was observed on average 9.5 times per month (range: 2-19 times/month). During focal samples, observers recorded activity state, social interactions, and vocalizations on a continuous basis (J Altmann, 1974). For social interactions, observers recorded the type of social behavior, the identity of the partner, and whether the interaction was initiated by the focal animal, the partner, or jointly. For vocalizations, observers recorded the type of call given, the identity of the partner, and whether the call was given by the focal animal or its partner. Encounters with humans and baboons from other troops were also recorded ad libitum (J Altmann, 1974). All behavioral data were collected on hand-held computers (Palm Zire 21) in the field and later transferred onto computers for error checking and storage in the NS Basic program. Adult and subadult dominance ranks were assessed by long-term UNBP observers each month based on decided agonistic contests and submissive behaviors.

### Fecal Collection, Hormonal Extraction, and Hormone Assays

We include a total of 562 fecal samples from the 31 mothers in this study, aiming to collect one sample per female each week (average=2.85 samples per mother per month). The protocol for collection, extraction, and storage have been validated and described in detail in primates (Jacinta C. Beehner & McCann, 2008). Within 10 minutes following deposition, the fecal sample was mixed thoroughly with a wooden spatula, and an aliquot of the mixed sample (∼ 0.5 g wet feces) was placed in 3 mL of a methanol/acetone solution (4:1). The solution was immediately homogenized using a battery-powered vortex. The weight of the dry fecal matter was later determined using a battery-powered, portable scale to ± 0.001 g. Approximately 4–8 h after sample collection, 2.5 mL of the fecal homogenate was filtered through a 0.2 μm polytetrafluoroethylene (PTFE) syringeless filter (Fisher cat #09-921-13), and the filter was then washed with an additional 0.7 mL of methanol/acetone (4:1). We then added 7 mL of distilled water to the filtered homogenate, capped and mixed the solution, and loaded it onto a reverse-phase C18 solid-phase extraction cartridge (Fisher cat #50-818-645). Prior to loading, Sep-Pak cartridges were prepped according to the manufacturer’s instructions (with 2 mL methanol followed by 5 mL distilled water). After the sample was loaded, the cartridge was washed with 2 mL of a sodium azide solution (0.1%). All samples were stored on cartridges in separate sealed bags containing silica beads. Cartridges were stored at ambient temperatures for up to 10 days, after which all samples were stored at subzero temperatures (− 20 °C) until transported to Arizona State University for analysis. In the laboratory, steroids were eluted from cartridges with 2.5 mL 100% methanol and subsequently stored at − 20 °C until the time of enzyme immunoassay (EIA).

We analyzed GCMs in our samples using a group-specific EIA for the measurement of immunoreactive 11β-hydroxyetiocholanolone (Frigerio, Dittami, Möstl, & Kotrschal, 2004), which has been used to monitor glucocorticoids in other primate species and validated biologically with an ACTH challenge test in olive baboons (e.g. Barbary macaque, Macaca sylvanus: (M. Heistermann, Palme, & Ganswindt, 2006; Shutt, Maclarnon, Heistermann, & Semple, 2007); Assamese macaque, Macaca assamensis: (Ostner, Heistermann, & Schülke, 2008); douc langur, Pygathrix nemaeus: (Michael Heistermann, Ademmer, & Kaumanns, 2004); Verraux’s sifaka, Propithecus verrauxi: (Fichtel, Kraus, Ganswindt, & Heistermann, 2007); olive baboons: *personal communication as cited in* Higham, MacLarnon, Heistermann, Ross, & Semple, 2009). We used assay 69a from Rupert Palme’s lab. The Palme lab provided 5ß-androstane- 3α,11b-di-ol-17-one-CMO-biotinyl-LC label, 5ß-androstane-3α,11b-di-ol-17-one- CMO:BSA antibody, and standard. Cross-reactivities for the 69a assay are characterized in: Ganswindt, Palme, Heistermann, Borragan, & Hodges, 2003.

We diluted baboon fecal extracts in assay buffer and used serial dilutions to compare the slope between the pooled samples and the assay standards. Slopes were not significantly different for the pooled baboon samples and the standard curve (F = 0.10, p=0.77). Samples were diluted 1:60 in assay buffer. The standards curve ranged from 3.9 to 250.0 pg/well. Samples were run in duplicate and CVs over 20% were eliminated (mean CV = 7.37%). We used low and high concentrations of pooled baboon samples as inter-assay controls on each plate. Inter-assay CVs were 18.6% and 24.4% respectively. Samples and standards were added to each plate in duplicate (50 uL/well), followed by 50 uL of biotin-labeled hormone and 50 uL of antibody to each well. Plates were incubated for at least 18 h at 4°C, and no more 24 hours. Plates were washed with a wash solution (PBS solution with 0.05% tween) and 150 uL of streptavidin-peroxidase was added to each well, incubated for one hour, and then the plate was washed again. We added 100 uL of TMB substrate solution to each well. Plates were incubated while shaking for 55-60 mins and the reaction was stopped with the addition of 50 uL of sulfuric acid and the plate was read at wavelength of 450 nn on a Synergy H2 plater reader.

### Data Analysis

#### Assessment of Mothers’ Early Life Adversity

We modified the cumulative early life adversity index used by the Amboseli Baboon Research Project (Rosenbaum et al., 2020; Tung et al., 2016; Weibel, Tung, Alberts, & Archie, 2020; Zipple et al., 2019) to fit our study population of olive baboons. We considered 5 measures to assess the adversity experienced by mothers in their early development. Three of these measures were also used in the Amboseli study: biomass during the birth year as an indicator of environmental conditions (the Amboseli Baboon Project used rainfall), group size at birth as an indicator of the extent of within-group competition, and early loss of mother.

A fourth measure, IBI, was also used in previous studies, but we interpreted the effect of IBI differently. In the Amboseli studies, researchers reasoned that shorter interbirth intervals following a female’s birth would indicate higher amounts of competition with a younger sibling. Short IBIs are also linked to increased mortality risk in macaques (Lee, Ruiz-Lambides, & Higham, 2019). However, longer IBIs might reflect poor maternal condition. In primates, both low rank and older age are associated with longer IBIs (reviewed in Harcourt, 1987; eg, baboons: Cheney et al., 2004; Smuts & Nicolson, 1989; chimpanzees: Roof, Hopkins, Izard, Hook, & Schapiro, 2005; gorillas: Robbins, Robbins, Gerald-Steklis, & Steklis, 2006; macaques: Ha, Robinette, & Sackett, 2000; Sugiyama & Ohsawa, 1982; Van Noordwijk & Van Schaik, 1999), and this is likely a result of poorer energy balance or greater social stress. The advent of *Opuntia stricta* in the diet lowered IBIs in this study population (UNBP unpublished data). Further, higher group size at birth is associated with longer interbirth intervals in our study population (Figure S1, Table S1). Thus, we consider longer IBIs to be an indicator of adversity in this population.

We added a fifth measure, primiparity to the early life adversity index because young, primiparous mothers must trade off investment in their own growth and their offspring’s growth, and have fewer bodily resources available during pregnancy and lactation (Jeanne Altmann & Alberts, 2005; Katie Hinde & Milligan, 2011; Pittet et al., 2017; Stearns, 1992; Wathes et al., 2007). The heightened energetic demands on primiparous mothers can result in negative outcomes for offspring such as lower birth weight (Setchell & Dixson, 2001) or increased mortality risk (Asian elephants: Mar, Lahdenperä, & Lummaa, 2012; howler monkeys: Glander, 1980; baboons: Smuts & Nicolson, 1989; vervets: L.A. Fairbanks & McGuire, 1995; but see macaques: Nuñez, Grote, Wechsler, Allen-Blevins, & Hinde, 2015). Thus, we consider primiparity to be a form of early life adversity.

Previous studies rely on binary scores for components of the early life adversity. We used continuous measures for all components of the early life adversity index except primiparity to avoid binning data, which reduces precision of information and requires arbitrary cutoffs. All of the continuous measures were normalized so values range from zero to one and can be summed to create a cumulative score. Primiparity was scored as 1 to indicate adversity for first born mothers, and 0 to indicate a lack of adversity for mothers who were not first born. All five scores were summed to create the cumulative adversity index.

Continuous measures:

a. Biomass: we used herbaceous biomass to determine drought years. We recorded monthly biomass data separately for two ranging areas. NMU troop occupied one ranging area and PHG, ENK, and YNT occupied the other range. Biomass was averaged for the year of each mother’s birth and this was reversed so less biomass was a higher adversity score.
b. Experienced group size: group size was defined as the number of adult and subadult males and females in the troop on the day the mother was born.
c. Maternal loss: maternal loss was defined as the age at which a female lost her own mother. This score was then inverted so that maternal loss at an earlier age is associated with a higher value. We include maternal loss after the period of nutritional independence because death of mother continues to have substantial effects on offspring survival and fitness even following weaning (Crockford et al., 2020; Foster et al., 2012; Nakamura et al., 2014; Samuni et al., 2020; Stanton et al., 2020). We use 4 years of age as a cutoff because we are interested in early life experiences and 4 years marks the earliest age at menarche in this population (Strum & Western, 1982). Mothers who lost their own mother after the age of 4 years received a zero for this component of early life adversity.
d. Maternal investment period: this was defined as the time between a female’s own birth to the birth of her next younger sibling. Here we consider longer investment periods to represent an adversity (as described above).

We also consider a cumulative adversity index with binary scores based on Tung et al. (2016) (methods described in the supplementary materials). We compare model fit of models based on binary and continuous indices, and report results in the supplementary materials (Tables S2-S5, Figures S2-S5).

We also consider the presence of *Opuntia stricta* in mothers’ early lives. Long-term UNBP observations show that animals in PHG, ENK, and YNT started to eat *O. stricta* fruit in 2000 regularly and animals in NMU started to eat it regularly in 2008. Based on these dates, we measured each mother’s age at introduction to *O. stricta:* year troop started to regularly consume *O. stricta* minus the year of mother’s birth. Age of zero is used if *O. stricta* was already present at birth.

### Measures from Mothers’ Current Reproductive Events in Adulthood

Maternal effort was calculated as the proportion of observation time spent nursing offspring and the proportion of time spent carrying offspring. Specifically, for each day of focal observation, we calculated the total amount of time that offspring spent nursing and being carried by their mothers, and divided this by the total number of minutes observed. We calculated maternal rank relative to the total number of females in the hierarchy (Levy et al., 2020), such that ranks range from 0 to 1 and higher numbers indicate better rank.

We consider two forms of current challenges that may influence maternal GCMs and protective behaviors. Humans pose a serious threat to baboons in the region. During this study period, we recorded 4 deaths (3 infants, 1 adult female) due to human-baboon conflict. Visits and immigration of unfamiliar males are associated with elevated glucocorticoid levels in chacma baboon mothers (J. C. Beehner, Bergman, Cheney, Seyfarth, & Whitten, 2005) and increased risk of wounding in olive baboons and gelada monkeys (*Theropithecus gelada*) (MacCormick et al., 2012; Schneider-Crease et al., 2020). Thus, we assessed current challenges as the sum of monthly encounters with humans and unfamiliar male baboons.

We treated offspring survival as a binary score: a score of one if the offspring died before two years old and a score of zero if the offspring survived to at least two year of age. Offspring who disappeared before reaching two years of age are assumed to have died and were scored as one. Offspring who survived to at least age 2 were scored as zero. Offspring who were alive but less than two years of age at the end of the study are excluded from the analysis because we do not know if they would have survived to the age of two years.

### Statistical Modeling

To determine what factors predicted GCM levels, we fit Gaussian models. To examine the probability of mortality before 2 years of age, we fit binomial models. To determine what factors predicted nursing and carrying time, we constructed zero-augmented gamma (ZAG) models. ZAG models are mixture models that combine a Bernoulli and gamma distribution. The Bernoulli component uses a logit link and estimates *p*, the probability of not observing the maternal behavior. The gamma component estimates the mean duration of maternal behavior, mu, and a shape parameter, *k*, given the duration > 0. Although the durational behaviors are proportions bound by zero and one, a gamma distribution is appropriate because the data are heavily skewed towards zero. The joint likelihood of duration of behavior is calculated by multiplying the likelihoods of the Bernoulli and gamma outcomes. Negative coefficients from the Bernoulli component indicate a lower probability of not observing the behavior, while positive values for the gamma component indicate higher durations of the behavior. The regression coefficients provide information about effects, but in the mixture models, it is challenging to interpret the joint effects on posterior predictions so we have included graphs of joint model predictions. We recommend focusing on the graphs of joint model predictions over the raw data. We fitted models using Hamilton Markov chain Monte Carlo (MCMC) with r-STAN v.2.18.2 (Stan Develiopment Team, 2018) in R v.3.3.2 (R Core Team, 2017) using the map2stan function in the ‘rethinking’ package v.1.59 (McElreath, 2016). Model code is available at: https://github.com/skpatter/Maternal_early_life_adversity.

In all models described below, we used weakly informative priors for our fixed effects, setting the mean to 0 and the standard deviation to 2. This method constrains parameter estimates to biologically plausible values, while allowing the information in the data to dominate information in the prior. We used non-centered parameterization for the varying effects, which helps the model sample more efficiently (McElreath, 2016). To verify that the models were insensitive to the chosen priors, we ran a series of models with both weakly informative priors and regularizing priors for all predictor parameters, and our results were unaffected. We used effective sample size and the Gelman-Rubin convergence diagnostic (Rhat) to evaluate the quality of our models.

We ran a set of models for each of the response variables: proportion of observed time spent nursing, proportion of observed time spent carrying, GCM levels, and mortality. To account for repeated measures of individuals, maternal ID (or offspring ID for the nursing and carrying models) is included as a varying effect. For the nursing, carrying, and GCM models, the predictor variables were early life adversity score of mother, current maternal relative rank, number of current monthly challenges, current monthly herbaceous biomass, mother’s age at introduction to *Opuntia stricta,* group size on day of observation, infant age on the day of observation, mother’s age on the day of observation, and infant sex with male as the reference category. GCM models included samples from pregnant and lactating females and “infant age” in this model ranges from -180 to 365 days. This age is squared in the GCM models because maternal GCM levels rise across pregnancy and decline following parturition (Jeanne Altmann, Lynch, Nguyen, Alberts, & Gesquiere, 2004; Jacinta C. Beehner, Nguyen, Wango, Alberts, & Altmann, 2006). For the mortality models, the predictor variables were early life adversity score of mother, current maternal relative rank, mother’s age at introduction to *O. stricta*, mother’s age at offspring’s birth, and group size at infant birth. The mortality model includes births prior to the study period, but we do not have access to data on monthly challenges and herbaceous biomass for this entire period, so these predictors are not included. Age at opuntia introduction was closely associated with troop membership. To avoid collinearity in predictor variables, we use only age at opuntia introduction in our models because we think this measure is more biologically meaningful. Additionally, we include group size to further account for variation among troops. All continuous predictor variables were standardized to a mean of zero and standard deviation of one.

For each output measure, we ran one model including rank and early life adversity and a second model including an interaction between rank and early life adversity. To compare model fits, we use WAIC (Widely Applicable Information Criterion) values. We use model averaging to plot the results from these two models. We ran each of these models with the cumulative early life adversity index and with individual measures of early life adversity. The fit of models with the cumulative index and individual measures are compared using WAIC scores. This results in 4 models per output measure: early life adversity index1 (rank and early life adversity), early life adversity index2 (rank* early life adversity), separate early life adversity 1 (rank and early life adversity), separate early life adversity 2 (rank* early life adversity). The cumulative early life adversity index models produced a better fit than models with separate early life adversity variables in 6 out 8 comparisons. We present results from the cumulative early life adversity index models below, and results from models with separate early life adversity variables in the Supplementary Materials (Table S6-S9 Figure S6-S9).

### Ethical Note

The study conformed to U.S. and Kenyan laws and was approved by the National Commission for Science and Technology of Kenya and the Kenya Wildlife Service. The project was approved by the Arizona State University Institutional Care and Use Committee.

## Results

### Early life adversity

Early life adversity scores ranged from 0.81 to 2.8 (out of 5) across mothers with a mean (and standard deviation) of 1.70 ± 0.57 (Figure 2). Mothers in NMU experienced the most adversity on average (mean=2.10, sd=0.45), followed by ENK (mean=1.44, sd=0.43), PHG (mean=1.37, sd=0.50), and YNT (mean=1.21, sd=0.18).

**Figure 2.**
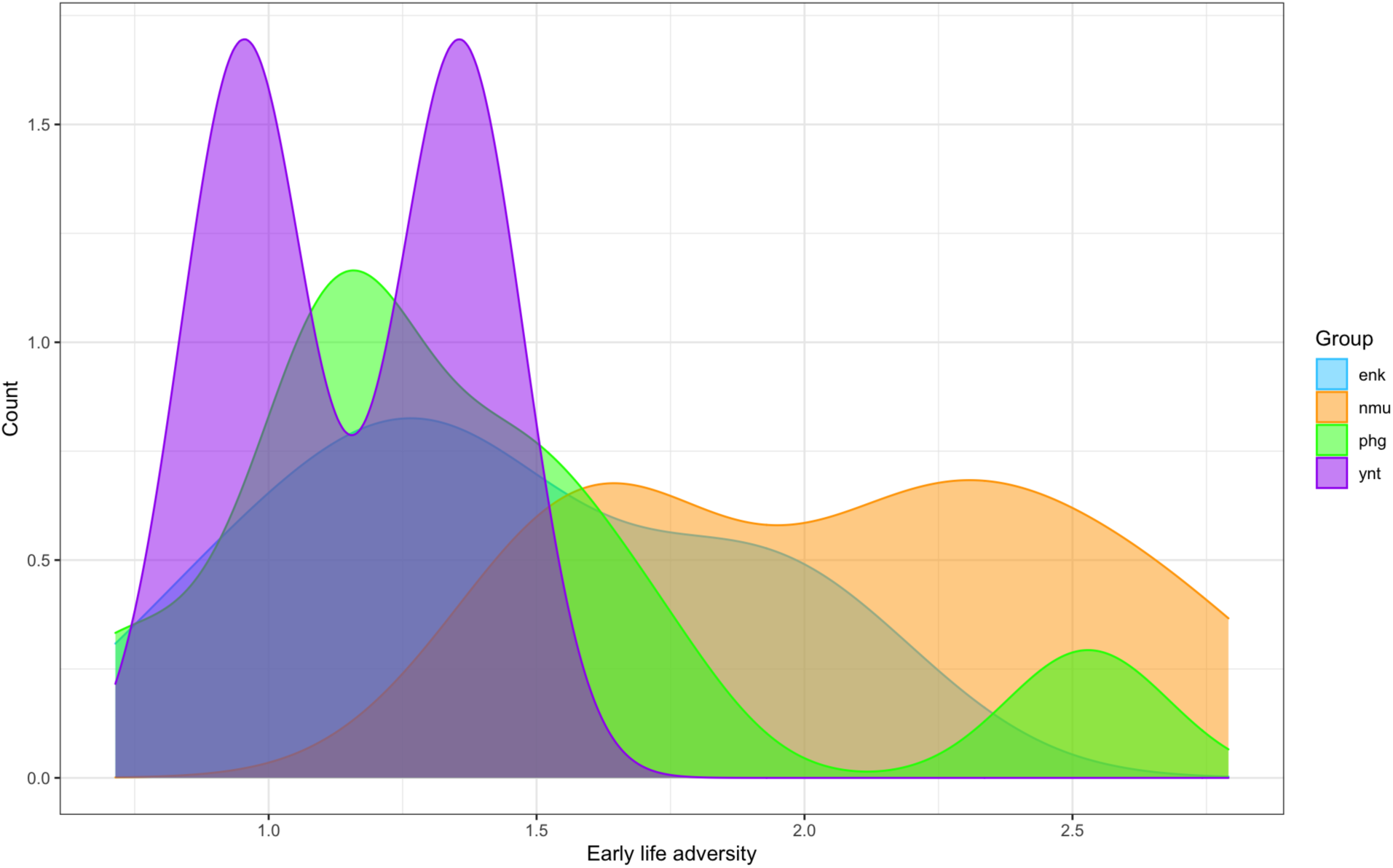
Distribution of early life adversity scores among females in the study

### Time spent nursing

Mothers who experienced more early life adversity nursed their offspring more than mothers who experienced less early life adversity (Figure 3, Table 2). Higher ranking mothers nursed their offspring less than lower ranking mothers. There was no interaction between early life adversity and rank, and the interaction between early life adversity and rank did not improve model fit (see WAIC scores and WAIC weights in Table 2; see interaction plot in Figure S10). Sons nursed more than daughters. Nursing time increased with the number of current monthly challenges, current monthly herbaceous biomass, and decreasing group size (Table 2).

**Figure 3.**
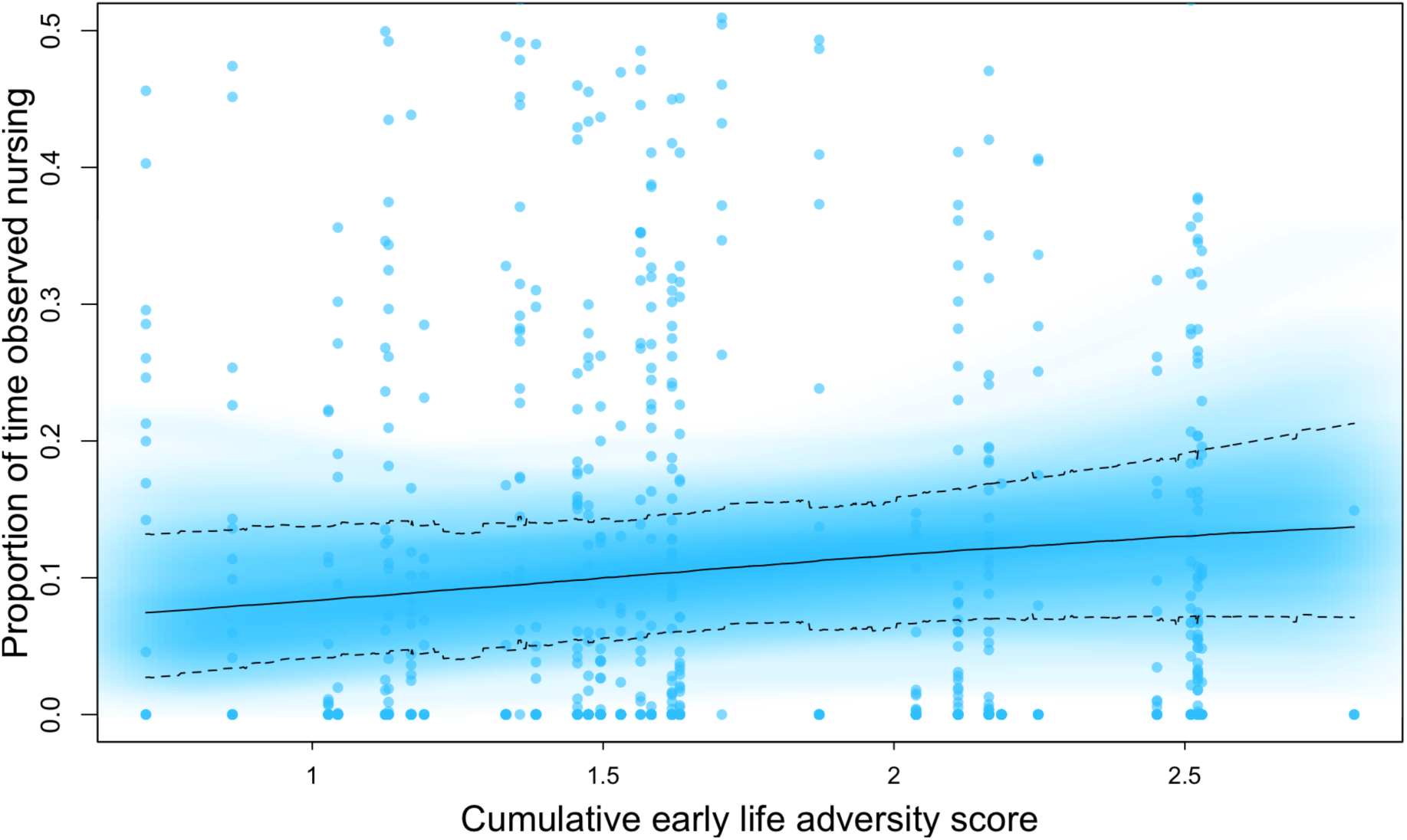
Nursing and early life adversity Model averaged posterior predictions for the influence of maternal cumulative early life adversity on the proportion of observation time offspring spent nursing. The solid line represents the mean estimate. The dashed lines represent the 89% highest posterior density interval. The blue cloud shows the full posterior predictions, with darker areas representing higher densities. Model sample sizes are as follows: 34 infants, 31 mothers, and 882 data points.

**Table 2.**
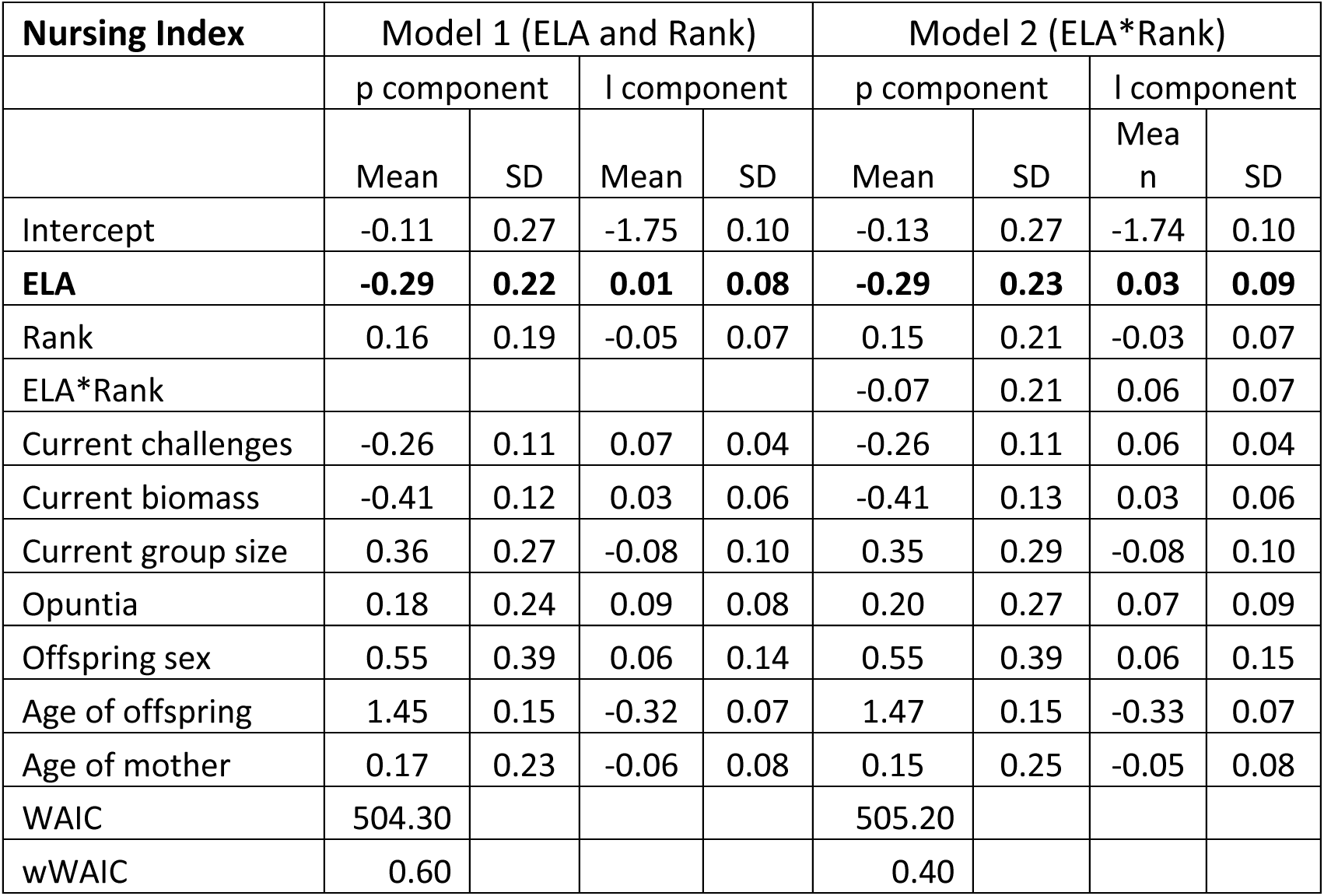
Coefficients for models evaluating the effect of maternal cumulative early life adversity (ELA) scores on offspring nursing time.

| Nursing Index | Model 1 (ELA and Rank) |  |  |  | Model 2 (ELA*Rank) |  |  |  |
| --- | --- | --- | --- | --- | --- | --- | --- | --- |
|  | p component |  | l component |  | p component |  | l component |  |
|  | Mean | SD | Mean | SD | Mean | SD | Mean | SD |
| Intercept | -0.11 | 0.27 | -1.75 | 0.10 | -0.13 | 0.27 | -1.74 | 0.10 |
| **ELA** | **-0.29** | **0.22** | **0.01** | **0.08** | **-0.29** | **0.23** | **0.03** | **0.09** |
| Rank | 0.16 | 0.19 | -0.05 | 0.07 | 0.15 | 0.21 | -0.03 | 0.07 |
| ELA*Rank |  |  |  |  | -0.07 | 0.21 | 0.06 | 0.07 |
| Current challenges | -0.26 | 0.11 | 0.07 | 0.04 | -0.26 | 0.11 | 0.06 | 0.04 |
| Current biomass | -0.41 | 0.12 | 0.03 | 0.06 | -0.41 | 0.13 | 0.03 | 0.06 |
| Current group size | 0.36 | 0.27 | -0.08 | 0.10 | 0.35 | 0.29 | -0.08 | 0.10 |
| Opuntia | 0.18 | 0.24 | 0.09 | 0.08 | 0.20 | 0.27 | 0.07 | 0.09 |
| Offspring sex | 0.55 | 0.39 | 0.06 | 0.14 | 0.55 | 0.39 | 0.06 | 0.15 |
| Age of offspring | 1.45 | 0.15 | -0.32 | 0.07 | 1.47 | 0.15 | -0.33 | 0.07 |
| Age of mother | 0.17 | 0.23 | -0.06 | 0.08 | 0.15 | 0.25 | -0.05 | 0.08 |
| WAIC | 504.30 |  |  |  | 505.20 |  |  |  |
| wWAIC | 0.60 |  |  |  | 0.40 |  |  |  |

### Time spent carrying

Mothers who experienced more early life adversity carried their offspring more than mothers who experienced less early life adversity (Figure 4, Table 3). Lower ranking mothers also carried their offspring more than higher ranking mothers. We did not find evidence for an interaction between rank and early life adversity. The model without an interaction between early life adversity and rank had a better fit than the model with this interaction (see WAIC in Table 3; see interaction plot in Figure S11). Sons were carried more than daughters. Carrying time increased with current monthly herbaceous biomass. Mothers who gained access to Opuntia later in their lives carried their offspring more than mothers who were born with access to the novel fruit (Table 3).

**Figure 4.**
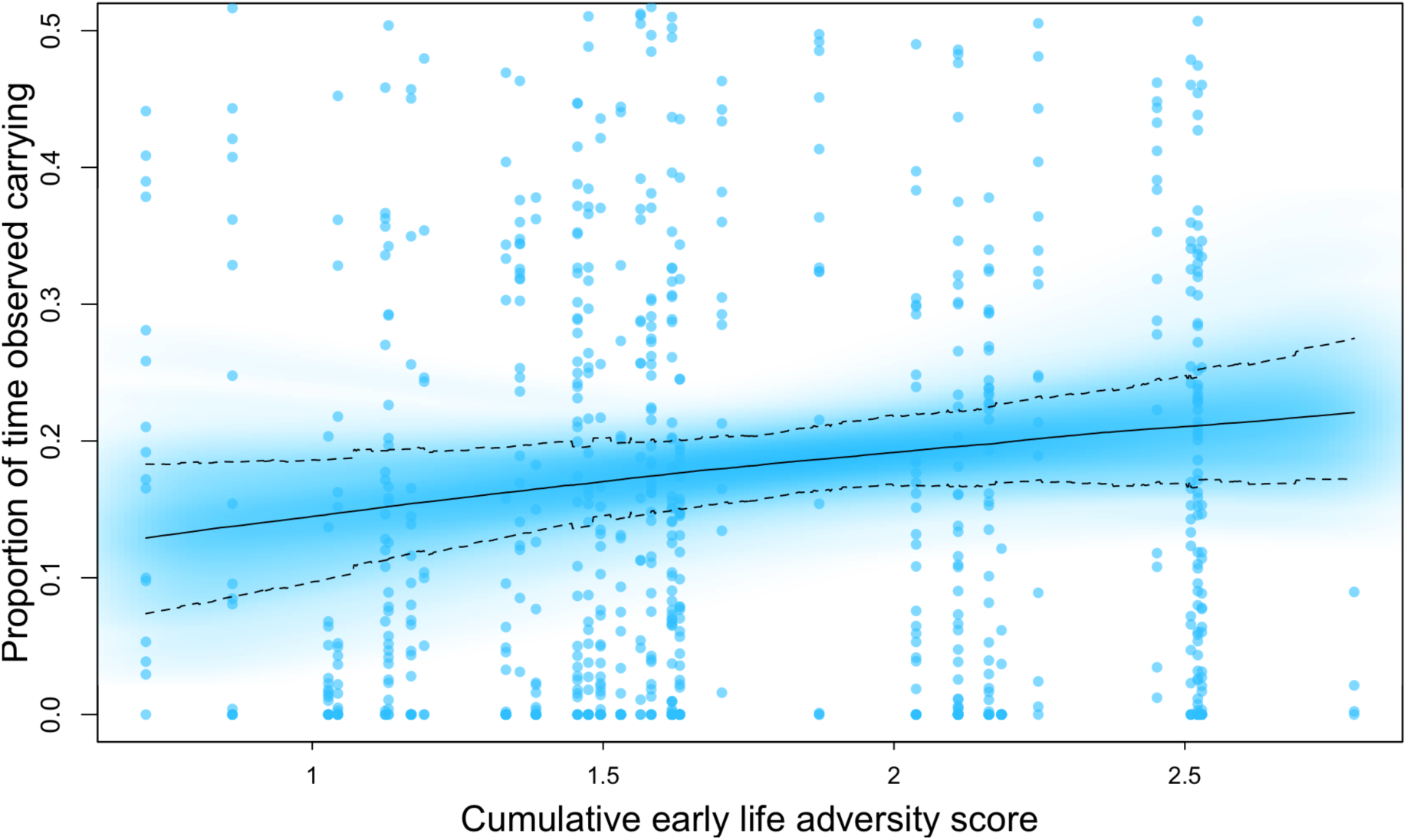
Carrying and early life adversity Model averaged posterior predictions for the influence of maternal cumulative early life adversity on the proportion of observation time spent carrying offspring. Model sample sizes are as follows: 34 infants, 31 mothers, and 882 data points.

**Table 3.** Coefficients for models evaluating the effect of maternal cumulative early life adversity (ELA) scores on offspring carrying time.

| Carrying Index | Model 1 (ELA and Rank) |  |  |  | Model 2 (ELA*Rank) |  |  |  |
| --- | --- | --- | --- | --- | --- | --- | --- | --- |
|  | p component |  | l component |  | p component |  | l component |  |
|  | Mean | SD | Mean | SD | Mean | SD | Mean | SD |
| Intercept | -1.54 | 0.28 | -1.62 | 0.07 | -1.56 | 0.29 | -1.63 | 0.07 |
| **ELA** | **-0.53** | **0.24** | **0.06** | **0.06** | **-0.53** | **0.24** | **0.04** | **0.06** |
| Rank | 0.37 | 0.21 | -0.05 | 0.05 | 0.37 | 0.22 | -0.06 | 0.05 |
| ELA*Rank |  |  |  |  | -0.02 | 0.21 | -0.03 | 0.05 |
| Current challenges | -0.08 | 0.14 | 0.01 | 0.03 | -0.08 | 0.14 | 0.01 | 0.03 |
| Current biomass | -0.02 | 0.14 | 0.06 | 0.04 | -0.02 | 0.14 | 0.06 | 0.04 |
| Current group size | -0.03 | 0.30 | 0.02 | 0.07 | -0.03 | 0.30 | 0.02 | 0.07 |
| Opuntia | 0.30 | 0.26 | -0.04 | 0.06 | 0.32 | 0.28 | -0.02 | 0.07 |
| Offspring sex | 0.22 | 0.41 | -0.04 | 0.10 | 0.23 | 0.42 | -0.04 | 0.10 |
| Age of offspring | 2.14 | 0.20 | -0.37 | 0.05 | 2.15 | 0.20 | -0.37 | 0.05 |
| Age of mother | -0.13 | 0.23 | 0.07 | 0.06 | -0.14 | 0.26 | 0.06 | 0.06 |
| WAIC | 29.20 |  |  |  | 31.20 |  |  |  |
| wWAIC | 0.73 |  |  |  | 0.27 |  |  |  |

### Maternal GCMs

Mothers who experienced more early life adversity had slightly higher GCM levels (Figure 5, Table 4). The nature of the relationship between early life adversity and GCMs did not differ across ranks, but the positive relationship was stronger among higher ranking mothers (see interaction plot Figure S12). The interaction between rank and early life adversity improved model fit (see WAIC in Table 4). Mothers of sons had higher GCMs than mothers of daughters. Maternal GCM levels decreased with more current monthly herbaceous biomass and higher current group size, and GCMs increased with maternal age (Table 4).

**Figure 5.**
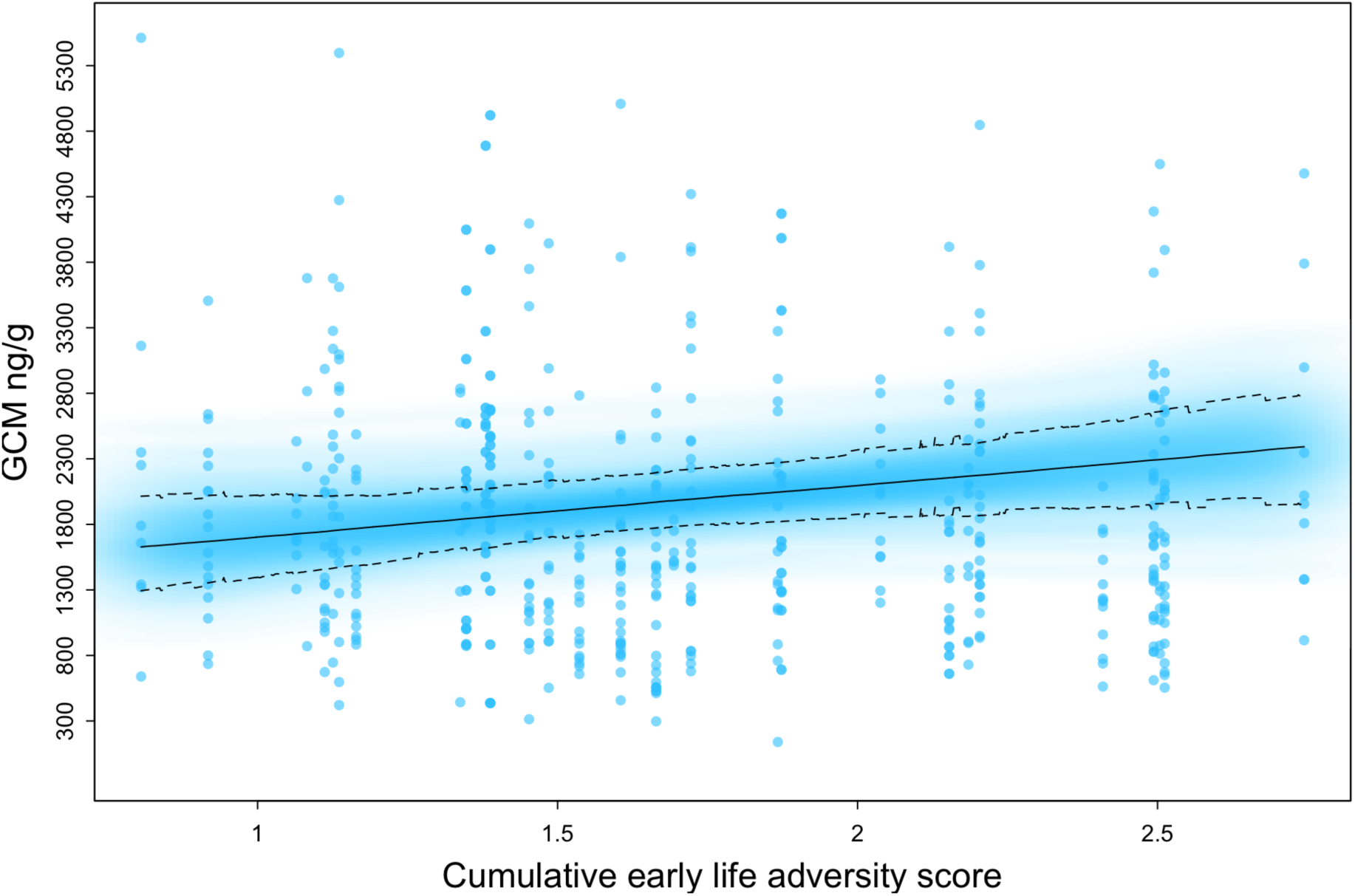
GCMs and early life adversity Model averaged posterior predictions for the influence of maternal cumulative early life adversity on adult GCM levels. Model sample sizes are as follows: 31 mothers and 562 data points.

**Table 4.** Coefficients for models evaluating the effect of maternal cumulative early life adversity (ELA) scores on adult GCM levels.

| GCM Index | Model 1<br>(ELA and Rank) |  | Model 2<br>(ELA*Rank) |  |
| --- | --- | --- | --- | --- |
|  | Mean | SD | Mean | SD |
| Intercept | 0.07 | 0.08 | 0.09 | 0.08 |
| **ELA** | **0.10** | **0.07** | **0.13** | **0.07** |
| Rank | 0.02 | 0.06 | 0.03 | 0.05 |
| ELA*Rank |  |  | 0.11 | 0.05 |
| Current challenges | -0.05 | 0.05 | -0.05 | 0.05 |
| Current biomass | -0.08 | 0.06 | -0.08 | 0.06 |
| Current group size | -0.17 | 0.09 | -0.18 | 0.08 |
| Opuntia | -0.05 | 0.07 | -0.08 | 0.07 |
| Offspring sex | -0.09 | 0.11 | -0.09 | 0.11 |
| Age of offspring | 0.05 | 0.05 | 0.04 | 0.05 |
| Age of offspring squared | -0.02 | 0.03 | -0.03 | 0.03 |
| Age of mother | 0.12 | 0.06 | 0.14 | 0.06 |
| WAIC | 1586.7 |  | 1589.0 |  |
| wWAIC | 0.24 |  | 0.76 |  |

### Offspring Mortality

Mothers who experienced more early life adversity gave birth to offspring with a higher probability of dying before 2 years of age than mothers who experienced less early life adversity, although there is considerable error (Figure 6, Table 5). Mothers’ dominance ranks did not predict offspring mortality and the interaction between early life adversity and rank did not improve the model fit (see WAIC in Table 5; see interaction plot in Figure S13). The probability of offspring mortality was higher among the groups with the smallest current group size and among older mothers (Table 5).

**Figure 6.**
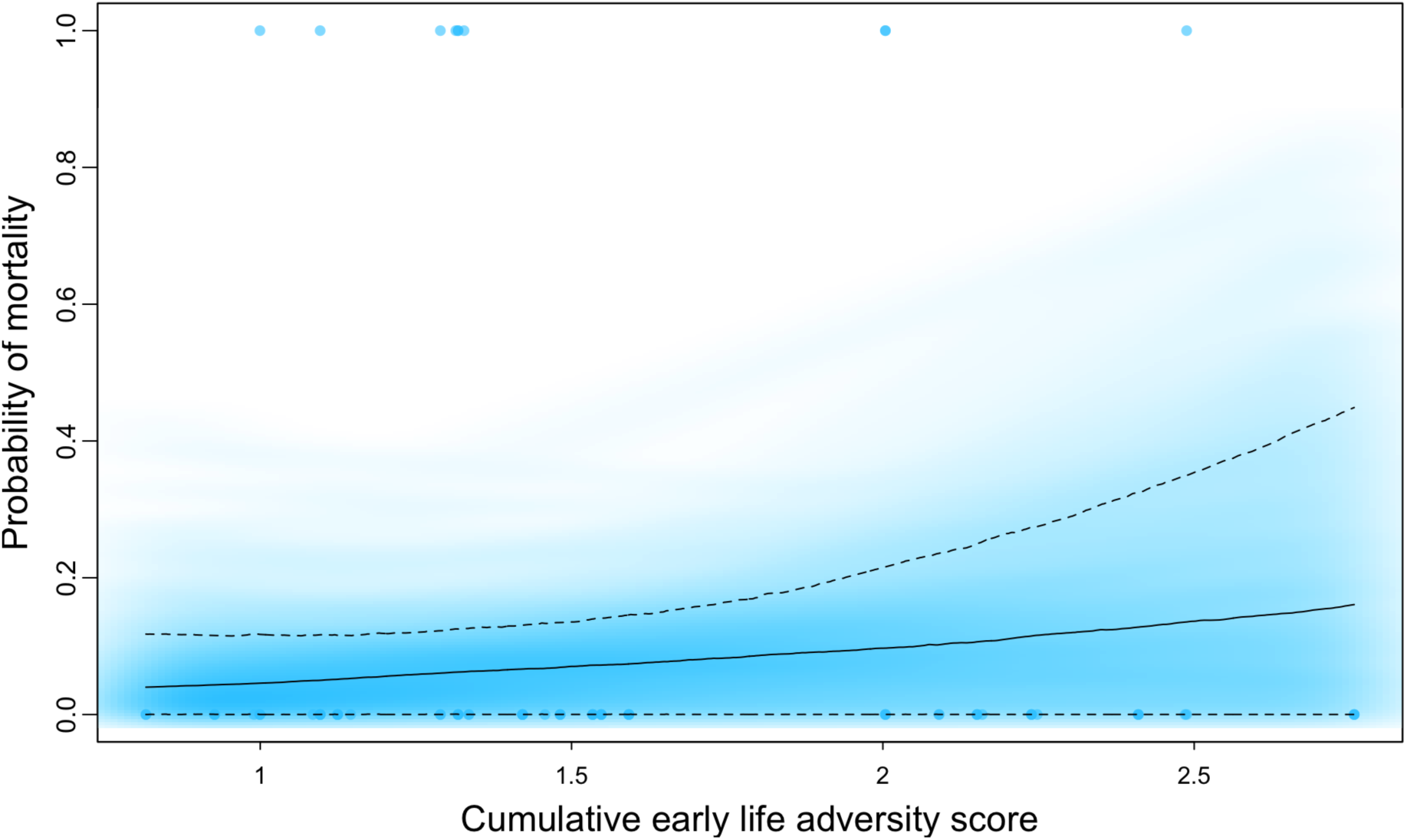
Offspring mortality and early life adversity Model averaged posterior predictions for the influence of maternal cumulative early life adversity on offspring mortality. Model sample sizes are as follows: 31 mothers and 80 data points.

**Table 5.**
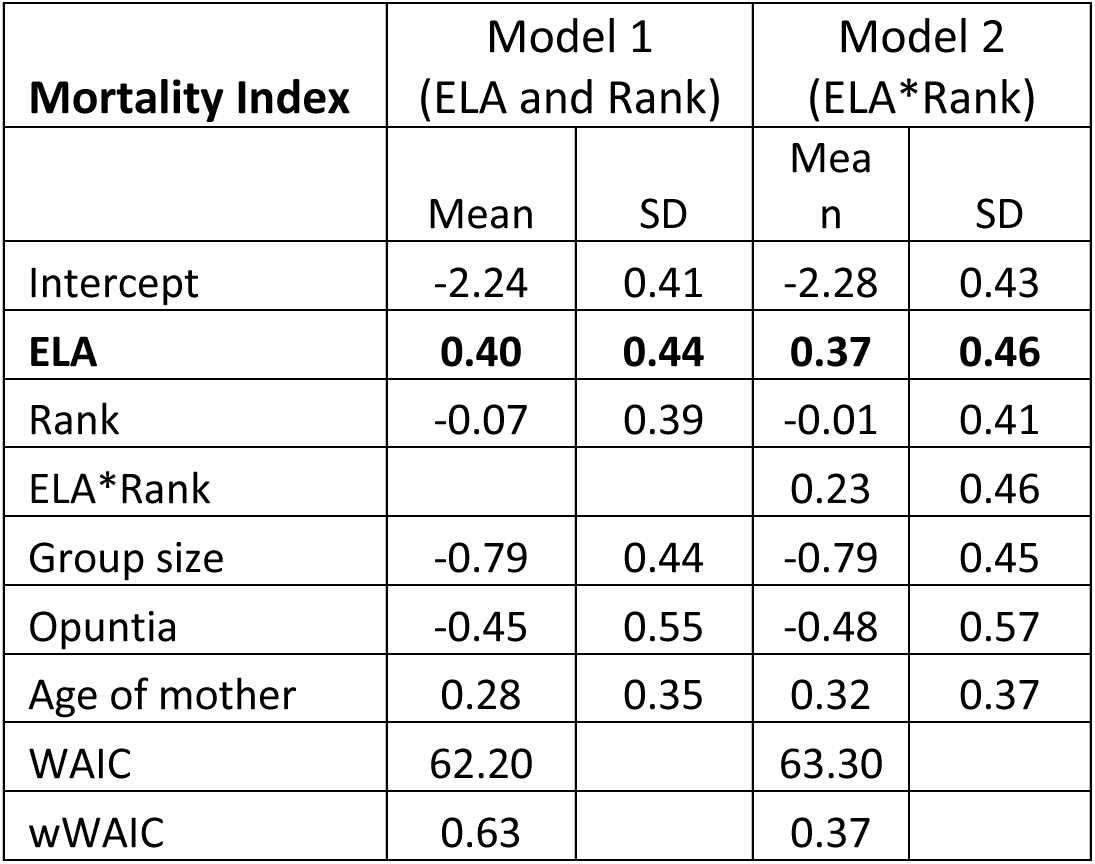
Coefficients for models evaluating the effect of maternal cumulative early life adversity (ELA) scores on offspring mortality.

## Discussion

Our findings substantiate that early life adversity constrains development with consequences for maternal effort, physiology, and offspring outcomes (Lea et al., 2015; Tung et al., 2016; Zipple et al., 2019). Mothers who experienced more early life adversity had higher concentrations of fecal glucocorticoid metabolites than did mothers with less early life adversity, and this was reflected in the behavior of mothers and their offspring. Mothers who experienced more early life adversity nursed and carried their offspring more than mothers who experienced less early life adversity. Greater maternal effort and elevated GCMs might be due to the poorer physical condition of mothers who experienced adversity. These patterns could also be due to social conditions. Female baboons with more early life adversity are less socially connected (Tung et al., 2016), so they might receive more aggression than females with less early life adversity and increase maternal protective behaviors (chimpanzees: Hemelrijk & De Kogel, 1989 ; rhesus macaques: Simpson & Howe, 1986; vervet monkeys: Fairbanks, 1996). Our findings also replicate results from muriquis, blue monkeys, and yellow baboons that linked maternal early life adversity to reduced offspring survival (Zipple et al., 2020, 2019). These observations add to a broader set of observations in plants, anthropods, fish, birds, and mammals that demonstrate the negative effects of early life adversity across generations (*reviewed in* Burton & Metcalfe, 2014).

In contrast to our predictions, high dominance rank did not buffer the effects of early life adversity. Other aspects of the social environment such as ties to close kin, social network position, or bonds with primary male associates might provide a buffer against the consequences associated with early life adversity and should be investigated in future work. For example, mountain gorillas who experience early maternal loss strengthen their social relationships, possibly in an effort to mitigate the consequences of maternal loss (Morrison, Eckardt, Colchero, Vecellio, & Stoinski, 2020). The buffering potential of sociality might be limited, however, as yellow baboons exposed to early life adversity experience weaker social bonds (Rosenbaum et al., 2020; Tung et al., 2016). Future studies in this study population on the links between early life adversity, sociality, and the outcomes tested here are needed to elucidate these patterns. Maternal dominance rank did influence patterns of maternal effort. Low-ranking mothers nursed and carried their offspring more than higher ranking mothers. These patterns may reflect the nutritional or social consequences of maternal rank. Other studies have also found that low ranking mothers nurse their offspring more (rhesus macaque daughters: Gomendio, 1989; yellow baboons: Nguyen, Gesquiere, Alberts, & Altmann, 2012) and carry their offspring more than higher ranking mothers (yellow baboons: Altmann, 1980; Altmann & Samuels, 1992; common marmosets: Digby, 1995 ; rhesus macaques: White & Hinde, 1975).

Maternal GCs are a key signal orchestrating offspring phenotype (Allen-Blevins et al., 2015; K. Hinde et al., 2015; Lu et al., 2019). Early life adversity has programming effects on neuroendocrine functioning and epigenetic changes to genes involved with HPA-axis regulation (Anacker et al., 2014; Maccari, Krugers, Morley-Fletcher, Szyf, & Brunton, 2014; Palma-Gudiel et al., 2015; Tyrka et al., 2016). The positive relationship between early life adversity and GCMs that we observed might be due to poorer physical condition of mothers who experienced adversity in their own early development, reduced social connectedness and heightened risk of aggression from conspecifics, or a combination of these physical and social mechanisms. Weak social bonds did not mediate the relationship between early life adversity and elevated GC concentrations in yellow baboons, suggesting social bonds might not play a major role in mediating effects of early life adversity on poor health outcomes in adulthood (Rosenbaum et al., 2020). Studies determining whether social bonds also play a minimal role in olive baboons and other systems are needed. Variation in GCM levels were not associated with maternal dominance rank. While some studies of primate females have found elevated GC levels among lower ranking individuals, most studies have not found a consistent relationship between rank and GCs (reviewed in Jacinta C. Beehner & Bergman, 2017; Carrera et al., 2020). Conceptually, rank serves as a proxy for condition insofar as access to resources and psychosocial stress are expected to vary, in part, as a function of social rank. Importantly, variation in environmental, group, and individual factors influence local resource competition and impact the extent of rank-mediated condition and may explain why rank effects are often absent.

Our analyses support the hypothesis that experiencing multiple adversities has compounding effects on adult outcomes (Hatch, 2005; Tung et al., 2016). In this study, we built models with a cumulative early life adversity index and models with each adverse condition considered separately. The cumulative index generally fit the data better than the individual measures, suggesting multiple adverse experiences compound in a biologically meaningful manner. However, examining the adverse measures separately also has its benefits. In the cumulative index models, high rank did not provide a buffer against early life adversity, but by examining measures separately, we found some aspects of early life adversity had larger effects on mortality, nursing, and carrying among lower ranking mothers, indicating high social status might act as a buffer against some forms of adversity.

Our early life adversity index differed from previous studies in several aspects. First, we interpreted the adverse effect of IBI differently. In the Amboseli yellow baboons, researchers treated shorter IBIs as an adversity because short IBIs indicate heightened competition with a younger sibling (e.g., Tung et al., 2016). However, we treated longer IBIs as an adversity because long IBIs might reflect poor maternal condition. In primates both low rank and older age are associated with longer IBIs (reviewed in Harcourt, 1987; eg, baboons: Cheney et al., 2004; Smuts & Nicolson, 1989; chimpanzees: Roof, Hopkins, Izard, Hook, & Schapiro, 2005; gorillas: Robbins, Robbins, Gerald-Steklis, & Steklis, 2006; macaques: Ha, Robinette, & Sackett, 2000; Sugiyama & Ohsawa, 1982; Van Noordwijk & Van Schaik, 1999), and this is likely a result of reduced access to food and poorer energy balance or greater social stress. In our study population, longer IBIs aligned with other forms of adversity as we would expect. The extent to which longer or shorter IBIs might be considered adverse likely varies by species, population, and conditions changing over time. Ultimately, there seems to be a U-shaped relationship with consequences arising from both the shortest and longest preceding and subsequent birth intervals (Conde-Agudelo, Rosas-Bermudez, Castaño, & Norton, 2012). Second, the cumulative early life adversity index that we constructed was based on normalized continuous scores of adversity, but analyses based on binary measures of adversity like those used by Tung et al (2016) produced a very similar pattern of results. One disadvantage of the binary index is that information is lost when continuous measures are treated as categorical, but a binary index might serve better if some measures do not have a linear effect or if extreme adverse situations drive the consequences of adversity. Decisions about adversity measures and indices should be determined for each species and population based on what is likely to be the most biologically relevant.

The current study has several limitations. While some conclusions can be drawn from the patterns of maternal effort and fecal GCMs established here, we lack important information on milk composition, quantity of milk transferred to offspring, and GC concentrations in milk. Further, it was beyond the scope of the present study to disentangle and differentiate the physical and social mechanisms linking mother’s early life adversity to her maternal effort and physiology. We were able to identify a link between a mother’s own early life adversity and her offspring’s mortality. However, due to our limited sample size, we were unable to directly link offspring survivorship to variation in maternal phenotype as a function of mother’s early life adversity. Our findings are consistent with the hypothesis that maternal effects play a role in the intergenerational transfer of early life adversity such that mothers’ own early life adversity influences their behavioral patterns and physiological signals during pregnancy and lactation, but shared genes and transgenerational epigenetics are also mechanisms that explain connections between maternal early experiences, phenotype, and offspring outcomes (Heard & Martienssen, 2014). Given our sample size and biomarkers assayed for the study population, we were unable to account for the role of genetics or epigenetics.

Research should aim to overcome the shortcomings of our current study. The amount of time spent nursing and carrying provide proxies for maternal effort, but data on maternal behavioral effort in conjunction with data on mother’s milk are needed to produce a comprehensive understanding of the complex, dynamic experiences of mothers and offspring. Future studies should also investigate the extent to which the patterns of maternal effort and physiology observed here are due to developmental constraints and/or social challenges. Research incorporating more detailed aspects of maternal care, male care, and the social environment in connection to multiple dimensions of offspring development and long-term offspring outcomes will continue to add important contributions to our understanding of maternal-offspring relations and developmental trajectories. We were limited by a small sample size and we were thus unable to directly test the impact of maternal effort and physiology on offspring outcomes, but such analyses are needed. While difficult to incorporate into wild primate studies, when possible, statistical analyses should use pedigrees to account for shared genes and estimate to what extent variance is explained by genetics and maternal effects (Brent, Ruiz-Lambides, & Platt, 2017; Kruuk, 2004; Wilson et al., 2010). Continued research on the health and fitness consequences of early life adversity, how social capital influences the effects of adversity, and mechanisms for persisting effects within and across generations will not only add to our understanding of variation in adult phenotype and infant development but might inform research on intervention practices in human health fields.

## Acknowledgements

We thank the Office of the President of the Republic of Kenya and the Kenya Wildlife Service for permission to conduct this field research. We thank Kate Abderholden, Megan Best, Megan Cole, Moira Donovan, Alexandra Duchesneau, Jessica Gunson, Molly McEntee, Laura Peña, Eila Roberts, and Leah Worthington for their contributions to data collection. We thank the staff of the Uaso Ngiro Baboon Project, particularly Jeremiah Lendira, James King’au, Joshua Lendira, and Frances Molo for their help and companionship in the field; David Muiruri for invaluable assistance with logistics and data management; and the African Conservation Centre for facilitating the UNBP project and assisting us with our work. We thank Jacinta Beehner, Sofia Carrera, Elizabeth Tinsley Johnson, Rupert Palme, Eila Roberts, and Sharmi Sen for guidance with hormonal analysis in the field and laboratory. We thank Joel Bray, Ian Gilby, Caitlin Hawley, Kevin Langergraber, Kevin Lee, Sarah Mathew, Tom Morgan, Shannon Roivas, India Schneider-Crease, and Veronika Städele for comments on an earlier draft of this paper. The research on which this paper was based was supported with funds to SKP from the National Science Foundation (BCS-1732172), the Leakey Foundation, Arizona State University, and with funds to JBS from Arizona State University. SKP was supported by the National Science Foundation Graduate Research Fellowship under grant no. 1841051. The funders had no role in study design, data collection and analysis, decision to publish, or preparation of the manuscript.

