## Supplementary Materials for "Effects of early life adversity on maternal effort and glucocorticoids in wild olive baboons"

Table S1. Interbirth interval as a function of group size and biomass at birth

| IBI Model | Mean | SD |
|---|---|---|
| Intercept | 0.00 | 0.14 |
| Number of adults & subadults | 0.42 | 0.14 |
| Biomass | 0.12 | 0.14 |

Figure S1. The relationship between IBI and group size at birth
Posterior predictions for the influence of the number of adult and subadult group members on length of interbirth intervals.

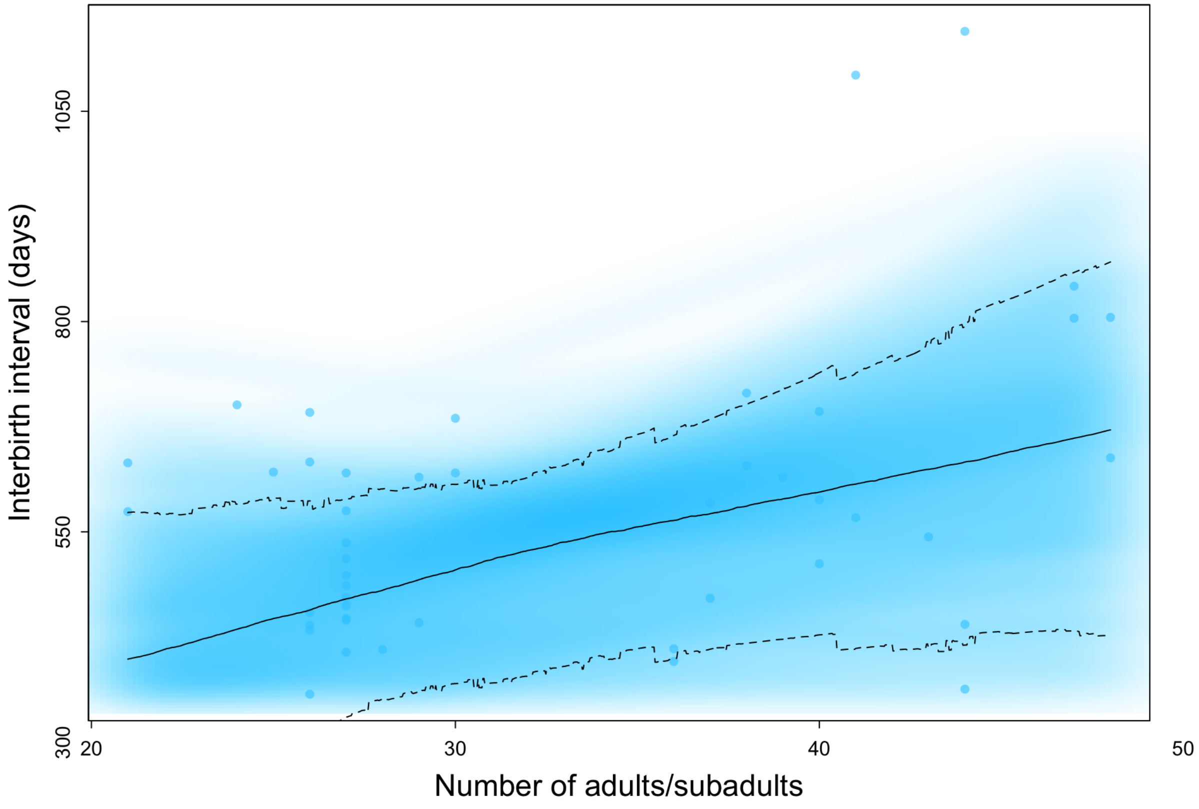


Methods for the early life adversity binary Index

The following 5 measures were scored as 0/1 and summed together to create a binary early life adversity index score

a) Biomass: We determined the mean and standard deviation of biomass for each ranging area from 1998-2012. Each range had a lower limit, which was ¾*standard deviation subtracted from the mean. If a mother was born in a year with biomass less than this lower limit, she was considered to have experienced this adversity.
b) Experienced group size: We calculated the upper quartile of densities based on data from all four troops 2008-2017. If a mother was born during a year when the density was greater than this upper quartile, she was considered to have experienced this adversity.
c) Maternal loss: Following Tung et al. (2016), a female is considered to have experienced this adversity if her own mother died before reaching 4 years of age.
d) Experienced interbirth interval: if the interval from a female's birth to her younger sibling's birth fell in the upper quartile of IBIs, she was considered to have experienced this adversity.
e) First born: if a female was her mother's first born, she experienced this adversity.

Table S2. Binary early life adversity (ELA) nursing models

| **Nursing Index Binary** | Model 1 (ELA and Rank) | | | | Model 2 (ELA*Rank) | | | |
|---|---|---|---|---|---|---|---|---|
| | p component | | l component | | p component | | l component | |
| | Mean | SD | Mean | SD | Mean | SD | Mean | SD |
| Intercept | -0.04 | 0.27 | -1.69 | 0.10 | -0.04 | 0.28 | -1.68 | 0.10 |
| **ELA Binary** | **-0.13** | **0.23** | **0.13** | **0.08** | **-0.12** | **0.24** | **0.13** | **0.08** |

| | | | | | | | | |
|---|---|---|---|---|---|---|---|---|
| Rank | 0.21 | 0.21 | -0.02 | 0.07 | 0.19 | 0.22 | -0.01 | 0.07 |
| ELA Binary*Rank | | | | | -0.11 | 0.22 | 0.04 | 0.07 |
| Current challenges | -0.26 | 0.11 | 0.06 | 0.04 | -0.26 | 0.11 | 0.06 | 0.04 |
| Current biomass | -0.37 | 0.12 | 0.04 | 0.06 | -0.37 | 0.12 | 0.05 | 0.06 |
| Current group size | 0.59 | 0.32 | -0.06 | 0.11 | 0.53 | 0.36 | -0.03 | 0.12 |
| Opuntia | -0.33 | 0.28 | -0.05 | 0.09 | -0.25 | 0.32 | -0.09 | 0.11 |
| Infant sex | 0.57 | 0.40 | -0.02 | 0.14 | 0.51 | 0.42 | 0.00 | 0.15 |
| Infant age | 1.39 | 0.15 | -0.35 | 0.07 | 1.41 | 0.15 | -0.35 | 0.07 |
| Mother's age | 0.40 | 0.21 | -0.05 | 0.07 | 0.37 | 0.23 | -0.04 | 0.07 |

**WAIC**
Model 1 (ELA and Rank):
Continuous ELA Index: WAIC = 501.4, weight = 0.81
Binary ELA Index: WAIC = 504.3, weight = 0.19
Model 2 (ELA*Rank):
Continuous ELA Index: WAIC = 503.1, weight = 0.74
Binary ELA Index: WAIC = 505.2, weight = 0.26

Figure S2. Binary early life adversity nursing models
Model averaged posterior predictions for the influence of maternal cumulative binary early life adversity on the proportion of observation time spent nursing.

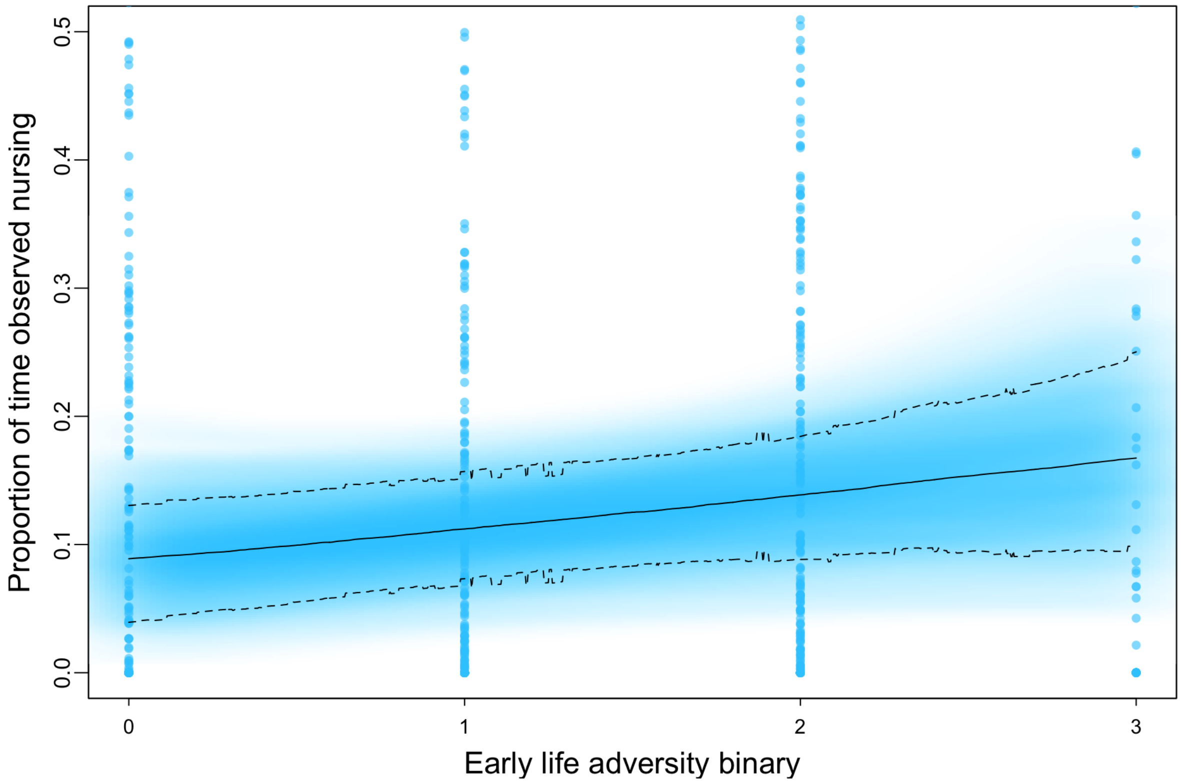


Table S3. Binary early life adversity (ELA) carrying models

| **Carrying Index Binary** | Model 1 (ELA and Rank) | | | | Model 2 (ELA*Rank) | | | |
|---|---|---|---|---|---|---|---|---|
| | p component | | l component | | p component | | l component | |
| | Mean | SD | Mean | SD | Mean | SD | Mean | SD |
| Intercept | -1.38 | 0.32 | -1.63 | 0.07 | -1.37 | 0.32 | -1.63 | 0.07 |
| **ELA Binary** | -0.16 | 0.28 | 0.05 | 0.06 | -0.23 | 0.28 | 0.05 | 0.06 |
| Rank | 0.41 | 0.24 | -0.04 | 0.05 | 0.46 | 0.23 | -0.04 | 0.05 |
| ELA Binary*Rank | | | | | 0.22 | 0.24 | 0.00 | 0.05 |
| Current challenges | -0.10 | 0.14 | 0.01 | 0.03 | -0.09 | 0.14 | 0.01 | 0.04 |
| Current biomass | 0.00 | 0.15 | 0.06 | 0.04 | 0.00 | 0.15 | 0.06 | 0.04 |
| Current group size | 0.13 | 0.37 | 0.02 | 0.08 | 0.29 | 0.44 | 0.02 | 0.09 |
| Opuntia | -0.27 | 0.35 | -0.01 | 0.07 | -0.43 | 0.40 | -0.02 | 0.08 |
| Infant sex | 0.10 | 0.48 | -0.04 | 0.10 | 0.21 | 0.50 | -0.04 | 0.11 |
| Infant age | 2.04 | 0.20 | -0.36 | 0.05 | 2.04 | 0.20 | -0.36 | 0.05 |
| Mother’s age | 0.16 | 0.24 | 0.04 | 0.05 | 0.21 | 0.27 | 0.04 | 0.06 |

**WAIC**

Model 1 (ELA and Rank):
Continuous ELA Index: WAIC = 29.2, weight = 0.75
Binary ELA Index: WAIC = 31.4, weight = 0.25
Model 2 (ELA*Rank):
Continuous ELA Index: WAIC = 21.2 weight = 0.82
Binary ELA Index: WAIC = 34.3, weight = 0.18

Figure S3. Binary early life adversity carrying models
Model averaged posterior predictions for the influence of maternal cumulative binary early life adversity on the proportion of observation time spent carrying offspring.

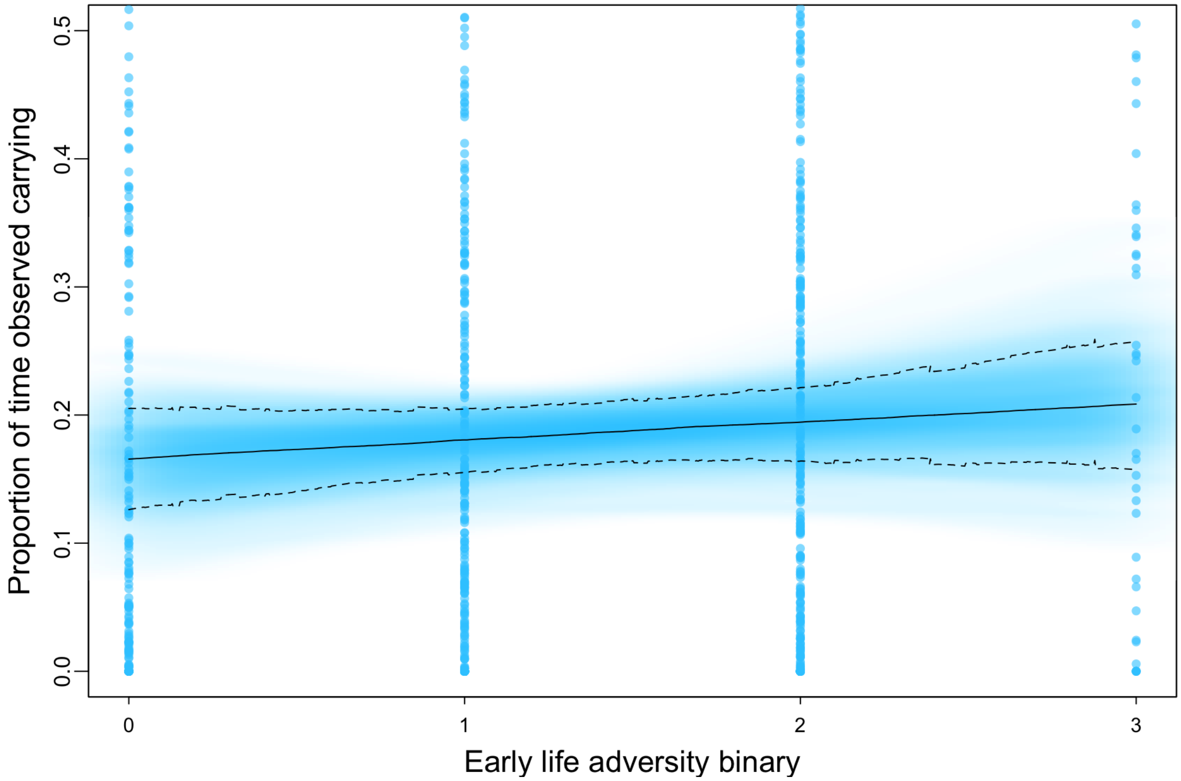


Table S4. Binary early life adversity (ELA) GCM models

| **GCM Index Binary** | Model 1 (ELA and Rank) | | Model 2 (ELA*Rank) | |
|---|---|---|---|---|
| | Mean | SD | Mean | SD |
| Intercept | 0.05 | 0.09 | 0.05 | 0.09 |
| **ELA binary** | **0.02** | **0.07** | **0.01** | **0.07** |
| Rank | 0.00 | 0.06 | 0.01 | 0.06 |

| | | | | |
|---|---|---|---|---|
| ELA binary*Rank | | | 0.07 | 0.06 |
| Current group size | -0.12 | 0.11 | -0.07 | 0.11 |
| Current challenges | -0.03 | 0.05 | -0.03 | 0.05 |
| Current biomass | -0.08 | 0.06 | -0.08 | 0.06 |
| Opuntia | -0.03 | 0.10 | -0.08 | 0.11 |
| Infant sex | -0.06 | 0.13 | -0.03 | 0.13 |
| Infant age | 0.07 | 0.05 | 0.07 | 0.05 |
| Infant age squared | -0.02 | 0.03 | -0.01 | 0.03 |
| Mother's age | 0.06 | 0.07 | 0.08 | 0.07 |

**WAIC**
Model 1 (ELA and Rank):
Continuous ELA Index: WAIC = 1589.0, weight = 1
Binary ELA Index: WAIC = 1604.7, weight = 0
Model 2 (ELA*Rank):
Continuous ELA Index: WAIC = 1586.0, weight = 1
Binary ELA Index: WAIC = 1604.7, weight = 0

Figure S4. Binary early life adversity GC models
Model averaged posterior predictions for the influence of maternal cumulative early life adversity on GC levels during pregnancy and lactation.

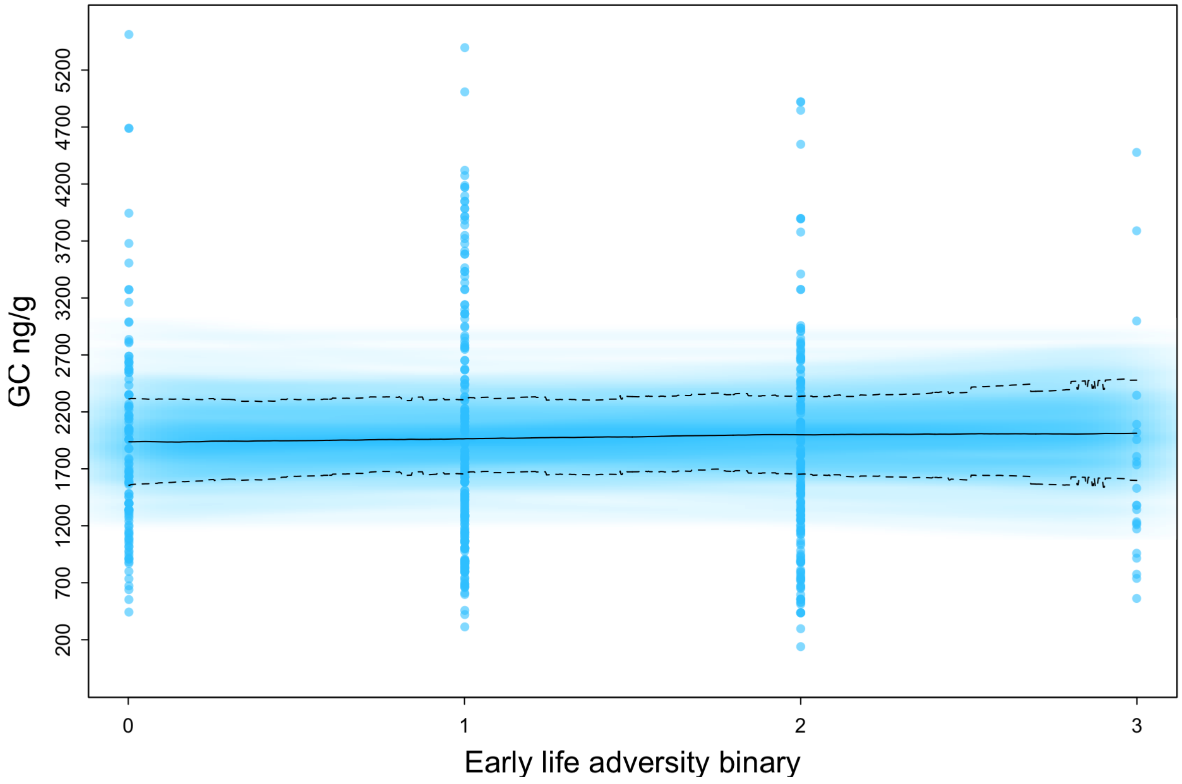

Table S5. Binary early life adversity (ELA) mortality models

| Mortality Index Binary | Model 1 (ELA and Rank) | | Model 2 (ELA*Rank) | |
|---|---|---|---|---|
| | Mean | SD | Mean | SD |
| Intercept | -2.23 | 0.43 | -2.29 | 0.43 |
| **ELA binary** | **0.49** | **0.44** | **0.59** | **0.46** |
| Rank | 0.01 | 0.39 | 0.04 | 0.40 |
| ELA binary*Rank | | | 0.48 | 0.44 |
| Current group size | -0.83 | 0.47 | -0.86 | 0.49 |
| Opuntia | -0.37 | 0.56 | -0.51 | 0.57 |
| Mother's age | 0.18 | 0.35 | 0.19 | 0.36 |

**WAIC**
Model 1 (ELA and Rank):
Continuous ELA Index: WAIC = 62.3, weight = 0.51
Binary ELA Index: WAIC = 62.4, weight = 0.49
Model 2 (ELA*Rank):
Continuous ELA Index: WAIC = 63.0, weight = 0.40
Binary ELA Index: WAIC = 62.2, weight = 0.60

Figure S5. Binary early life adversity mortality models
Model averaged posterior predictions for the influence of maternal cumulative binary early life adversity on the probability of offspring mortality.

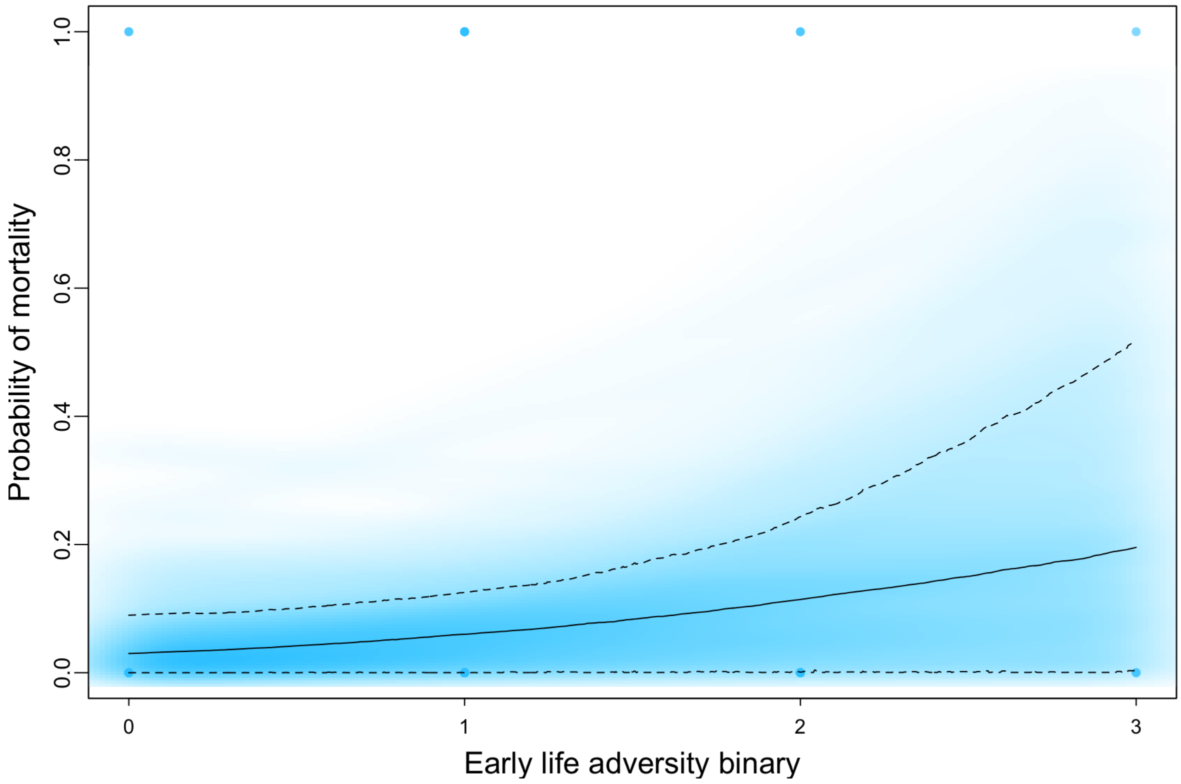


Table S6. Nursing models with separate early life adversity (ELA) measures

| **Nursing Separate** | Model 1 (ELA and Rank) | | | | Model 2 (ELA*Rank) | | | |
|---|---|---|---|---|---|---|---|---|
| | p component | | l component | | p component | | l component | |
| | Mean | SD | Mean | SD | Mean | SD | Mean | SD |
| Intercept | 0.02 | 0.30 | -1.67 | 0.10 | -0.14 | 0.39 | -1.58 | 0.12 |
| **Group size ELA** | **-0.24** | **0.45** | **0.20** | **0.14** | **-0.06** | **0.52** | **0.04** | **0.15** |
| **Biomass ELA** | **0.31** | **0.27** | **-0.11** | **0.09** | **0.35** | **0.30** | **-0.09** | **0.09** |
| **IBI ELA** | **-0.21** | **0.25** | **0.02** | **0.07** | **-0.18** | **0.28** | **0.00** | **0.07** |
| **Maternal loss ELA** | **0.23** | **0.34** | **-0.08** | **0.11** | **0.71** | **0.88** | **-0.34** | **0.30** |
| **Parity ELA** | **-0.37** | **0.57** | **-0.30** | **0.18** | **-0.17** | **0.74** | **-0.18** | **0.24** |
| Rank | 0.22 | 0.23 | -0.08 | 0.07 | 0.37 | 0.39 | -0.24 | 0.13 |
| Rank*group size | | | | | -0.10 | 0.24 | -0.06 | 0.06 |
| Rank*drought | | | | | -0.20 | 0.22 | 0.08 | 0.06 |
| Rank*ibi | | | | | -0.15 | 0.32 | 0.04 | 0.08 |
| Rank*maternal loss | | | | | -0.64 | 1.08 | 0.35 | 0.39 |
| Rank*parity | | | | | 0.31 | 0.76 | 0.30 | 0.24 |
| Current challenges | -0.24 | 0.11 | 0.07 | 0.04 | -0.24 | 0.11 | 0.07 | 0.04 |

| | | | | | | | | |
|---|---|---|---|---|---|---|---|---|
| Current biomass | -0.34 | 0.12 | 0.03 | 0.06 | -0.34 | 0.13 | 0.05 | 0.06 |
| Current group size | 0.98 | 0.51 | -0.21 | 0.15 | 0.88 | 0.58 | -0.02 | 0.16 |
| Opuntia | -0.55 | 0.38 | 0.07 | 0.11 | -0.49 | 0.45 | -0.07 | 0.12 |
| Infant sex | 0.64 | 0.42 | 0.07 | 0.13 | 0.69 | 0.48 | 0.07 | 0.13 |
| Infant age | 1.44 | 0.15 | -0.33 | 0.06 | 1.46 | 0.15 | -0.32 | 0.06 |
| Mother's age | 0.23 | 0.3 | 0.14 | 0.09 | 0.33 | 0.36 | 0.02 | 0.10 |
| WAIC | 505.5 | | | | 509.9 | | | |
| wWAIC | 0.90 | | | | 0.10 | | | |

**WAIC**

Model 1 (ELA and Rank):
Cumulative ELA Index: WAIC = 504.3, weight = 0.64
Separate ELA measures: WAIC = 505.5, weight = 0.36
Model 2 (ELA*Rank):
Cumulative ELA Index: WAIC = 505.2 weight = 0.91
Separate ELA measures: WAIC = 509.9, weight = 0.09

Figure S6. Relationship between nursing and each separate early life adversity measure
Model averaged posterior predictions for the influence of each maternal early life adversity measure on the proportion of observation time spent nursing for lower and higher ranking mothers.

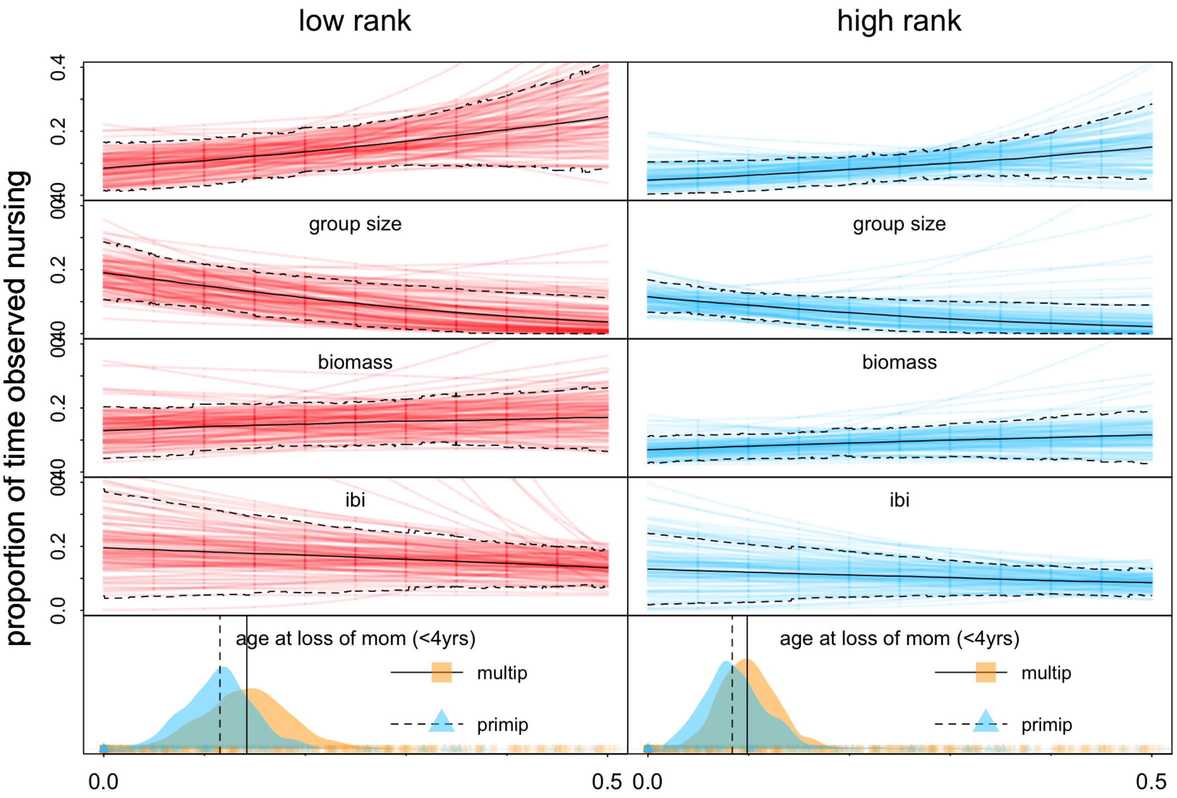


Table S7. Carrying models with separate early life adversity (ELA) measures

| **Carrying Separate** | Model 1 (ELA and Rank) | | | | Model 2 (ELA*Rank) | | | |
|---|---|---|---|---|---|---|---|---|
| | p component | | l component | | p component | | l component | |
| | Mean | SD | Mean | SD | Mean | SD | Mean | SD |
| Intercept | -1.22 | 0.35 | -1.62 | 0.08 | -1.29 | 0.45 | -1.51 | 0.11 |
| **Group size ELA** | **0.32** | **0.50** | **0.07** | **0.11** | **0.43** | **0.58** | **-0.03** | **0.14** |
| **Biomass ELA** | **-0.33** | **0.30** | **0.11** | **0.07** | **-0.33** | **0.36** | **0.07** | **0.08** |
| **IBI ELA** | **-0.33** | **0.28** | **0.04** | **0.05** | **-0.30** | **0.31** | **0.01** | **0.06** |
| **Maternal loss ELA** | **-0.19** | **0.36** | **0.14** | **0.08** | **-0.68** | **0.91** | **0.50** | **0.28** |
| **First born ELA** | **-1.05** | **0.61** | **0.07** | **0.13** | **-1.00** | **0.81** | **-0.02** | **0.19** |
| rank | **0.32** | **0.24** | **-0.06** | **0.05** | **0.48** | **0.42** | **-0.21** | **0.12** |
| rank*group size | | | | | -0.16 | 0.28 | 0.01 | 0.05 |
| rank*biomass | | | | | 0.04 | 0.26 | -0.07 | 0.05 |
| rank*ibi | | | | | 0.06 | 0.38 | -0.04 | 0.08 |
| rank*maternal loss | | | | | 0.63 | 1.11 | -0.51 | 0.36 |
| rank*first born | | | | | 0.04 | 0.82 | -0.07 | 0.19 |
| Current challenges | -0.03 | 0.14 | 0.00 | 0.03 | -0.03 | 0.15 | 0.01 | 0.04 |

| | | | | | | | | |
|---|---|---|---|---|---|---|---|---|
| Current biomass | 0.03 | 0.15 | 0.06 | 0.04 | 0.03 | 0.16 | 0.07 | 0.05 |
| Current group size | 0.23 | 0.57 | -0.09 | 0.12 | 0.20 | 0.66 | 0.00 | 0.15 |
| Opuntia | -0.50 | 0.44 | 0.08 | 0.09 | -0.49 | 0.54 | 0.03 | 0.11 |
| Infant sex | 0.26 | 0.52 | -0.08 | 0.11 | 0.16 | 0.62 | -0.06 | 0.12 |
| Infant age | 2.14 | 0.21 | -0.37 | 0.05 | 2.20 | 0.21 | -0.37 | 0.05 |
| Mother's age | 0.30 | 0.35 | 0.09 | 0.07 | 0.34 | 0.42 | 0.04 | 0.09 |
| WAIC | 34.6 | | | | 40.1 | | | |
| wWAIC | 0.94 | | | | 0.06 | | | |

**WAIC**

Model 1 (ELA and Rank):
Cumulative ELA Index: WAIC = 29.2, weight = 0.94
Separate ELA measures: WAIC = 34.6, weight = 0.06
Model 2 (ELA*Rank):
Cumulative ELA Index: WAIC = 31.2 weight = 0.99
Separate ELA measures: WAIC = 40.1, weight = 0.01

Figure S7. Relationship between carrying and each separate early life adversity measure
Model averaged posterior predictions for the influence of each maternal early life adversity measure on the proportion of observation time spent carrying offspring for lower and higher ranking mothers.

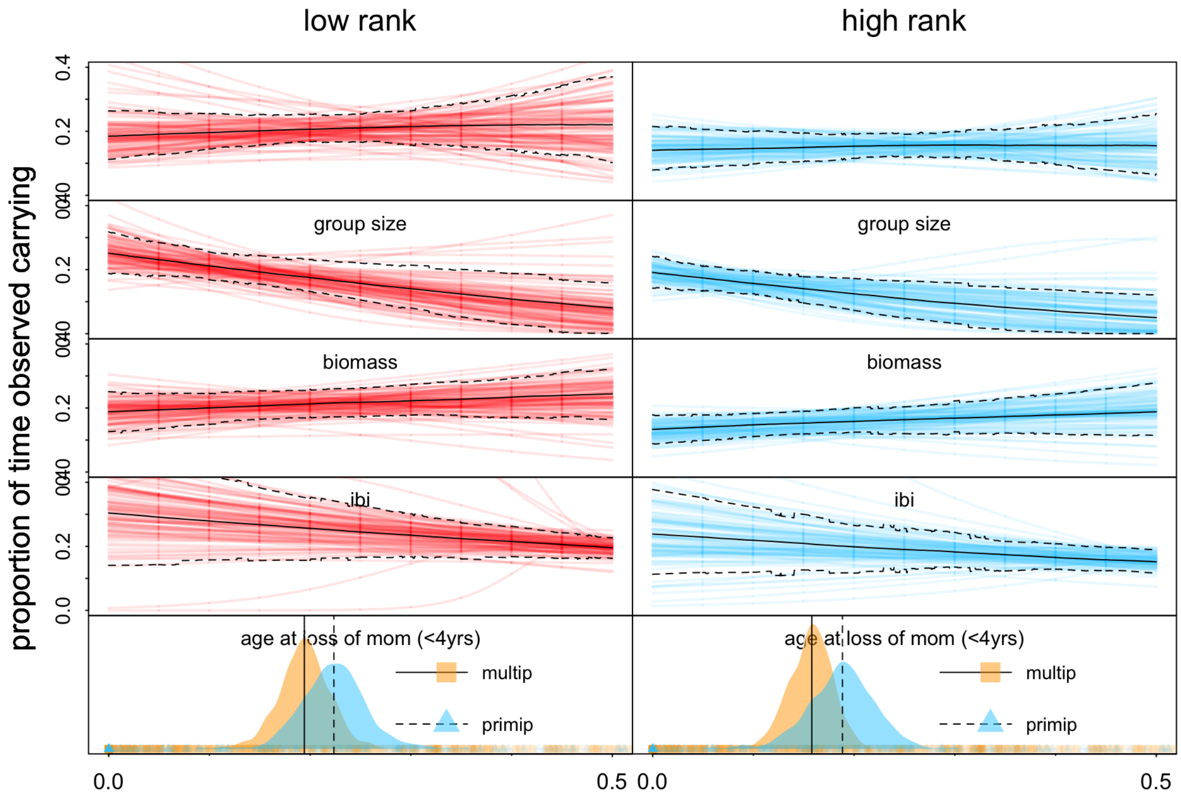


Table S8. GCM models with separate early life adversity (ELA) measures

| **GCM Separate** | Model 1 (ELA and Rank) | | Model 2 (ELA*Rank) | |
|---|---|---|---|---|
| | Mean | SD | Mean | SD |
| Intercept | 0.02 | 0.08 | 0.05 | 0.09 |
| **Group size ELA** | **0.32** | **0.12** | **0.33** | **0.16** |
| **Biomass ELA** | **-0.02** | **0.08** | **-0.06** | **0.09** |
| **IBI ELA** | **-0.02** | **0.06** | **-0.03** | **0.07** |
| **Maternal loss ELA** | **-0.04** | **0.06** | **-0.21** | **0.20** |
| **Parity ELA** | **0.12** | **0.16** | **0.28** | **0.29** |
| Rank | -0.02 | 0.06 | -0.07 | 0.09 |
| Group size*rank | | | 0.01 | 0.08 |
| Biomass*rank | | | 0.00 | 0.07 |
| IBI*rank | | | 0.11 | 0.09 |
| Maternal loss*rank | | | 0.19 | 0.26 |
| Parity*rank | | | 0.18 | 0.28 |

| | | | | |
|---|---|---|---|---|
| Opuntia | -0.05 | 0.13 | -0.15 | 0.15 |
| Current group size | -0.35 | 0.14 | -0.31 | 0.15 |
| Current challenges | -0.04 | 0.05 | -0.05 | 0.05 |
| Current biomass | -0.09 | 0.06 | -0.08 | 0.06 |
| Infant sex | -0.04 | 0.12 | -0.04 | 0.12 |
| Infant age | 0.06 | 0.05 | 0.06 | 0.06 |
| Infant age squared | -0.02 | 0.03 | -0.02 | 0.03 |
| Mother’s age | 0.16 | 0.06 | 0.17 | 0.06 |
| WAIC | 1587.2 | | 1592.2 | |
| wWAIC | 0.92 | | 0.08 | |

**WAIC**
Model 1 (ELA and Rank):
Cumulative ELA Index: WAIC = 1588.7, weight = 0.32
Separate ELA measures: WAIC = 1587.2, weight = 0.68
Model 2 (ELA*Rank):
Cumulative ELA Index: WAIC = 1586.0, weight = 0.96
Separate ELA measures: WAIC = 1592.2, weight = 0.04

Figure S8. Relationship between GCs and each separate early life adversity measure
Model averaged posterior predictions for the influence of each maternal early life adversity measure on GC levels for lower and higher ranking mothers.

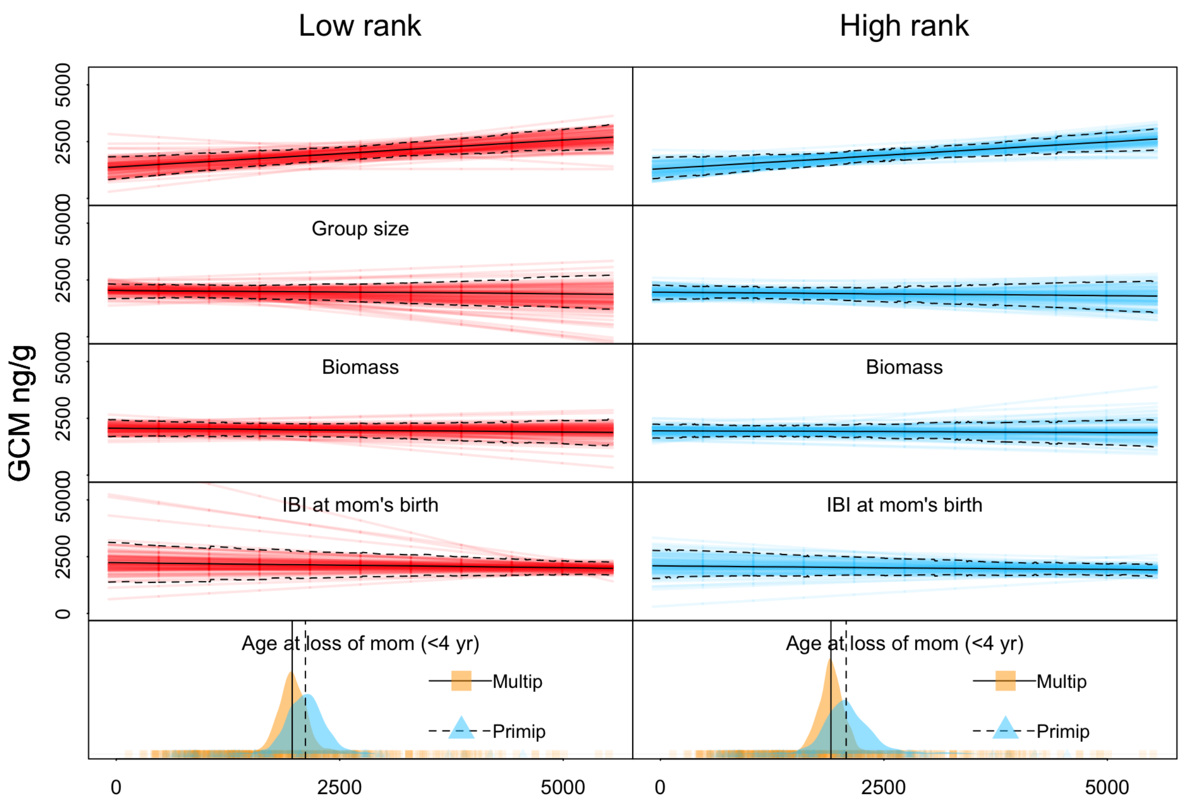


Table S9. Infant mortality models with separate early life adversity (ELA) measures

| **Mortality Separate** | Model 1 (ELA and Rank) | | Model 2 (ELA*Rank) | |
|---|---|---|---|---|
| | Mean | SD | Mean | SD |
| Intercept | -2.36 | 0.49 | -2.43 | 0.51 |
| **Group size ELA** | **-0.11** | **0.62** | **-0.07** | **0.69** |
| **Biomass ELA** | **-0.52** | **0.58** | **-0.61** | **0.57** |
| **IBI ELA** | **0.32** | **0.47** | **0.26** | **0.53** |
| **Maternal loss ELA** | **-0.12** | **0.47** | **-0.72** | **0.60** |
| **Parity ELA** | **-0.27** | **0.81** | **-0.14** | **0.82** |
| Rank | 0.04 | 0.45 | -0.16 | 0.53 |
| Group size*rank | | | -0.43 | 0.54 |
| Biomass*rank | | | 0.05 | 0.54 |
| IBI*rank | | | 0.98 | 0.56 |
| Maternal loss*rank | | | 1.05 | 0.79 |
| Parity*rank | | | 0.21 | 0.80 |
| Current group size | -0.82 | 0.52 | -0.80 | 0.53 |

| | | | | |
|---|---|---|---|---|
| Opuntia | -0.15 | 0.66 | -0.40 | 0.72 |
| Mother's age | 0.29 | 0.41 | 0.26 | 0.42 |
| WAIC | 65.7 | | 60.2 | |
| wWAIC | 0.06 | | 0.94 | |

**WAIC**
Model 1 (ELA and Rank):
Cumulative ELA Index: WAIC = 62.3, weight = 0.84
Separate ELA measures: WAIC = 65.7, weight = 0.16
Model 2 (ELA*Rank):
Cumulative ELA Index: WAIC = 63.0, weight = 0.20
Separate ELA measures: WAIC = 60.2, weight = 0.80

Figure S9. Relationship between infant mortality and each separate early life adversity measure
Model averaged posterior predictions for the influence of each maternal early life adversity measure on the probability of offspring mortality for lower and higher ranking mothers.

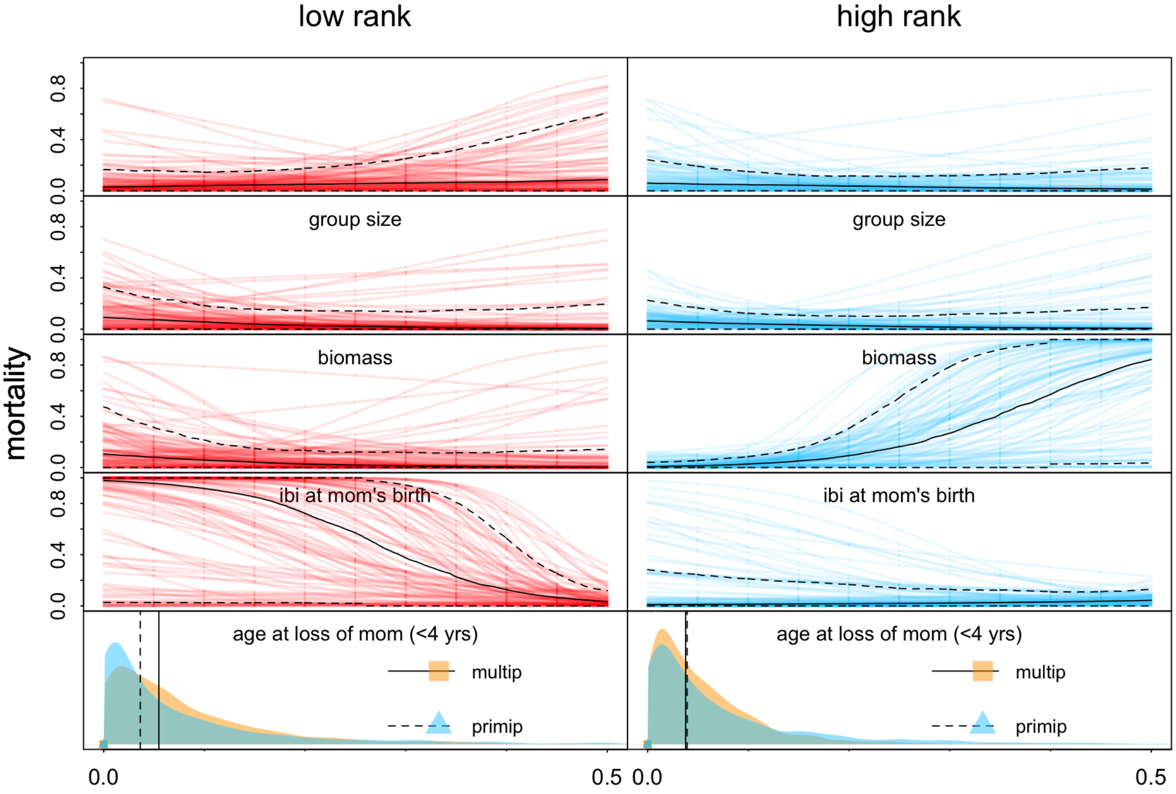

Figure S10. Nursing and early life adversity by maternal dominance rank

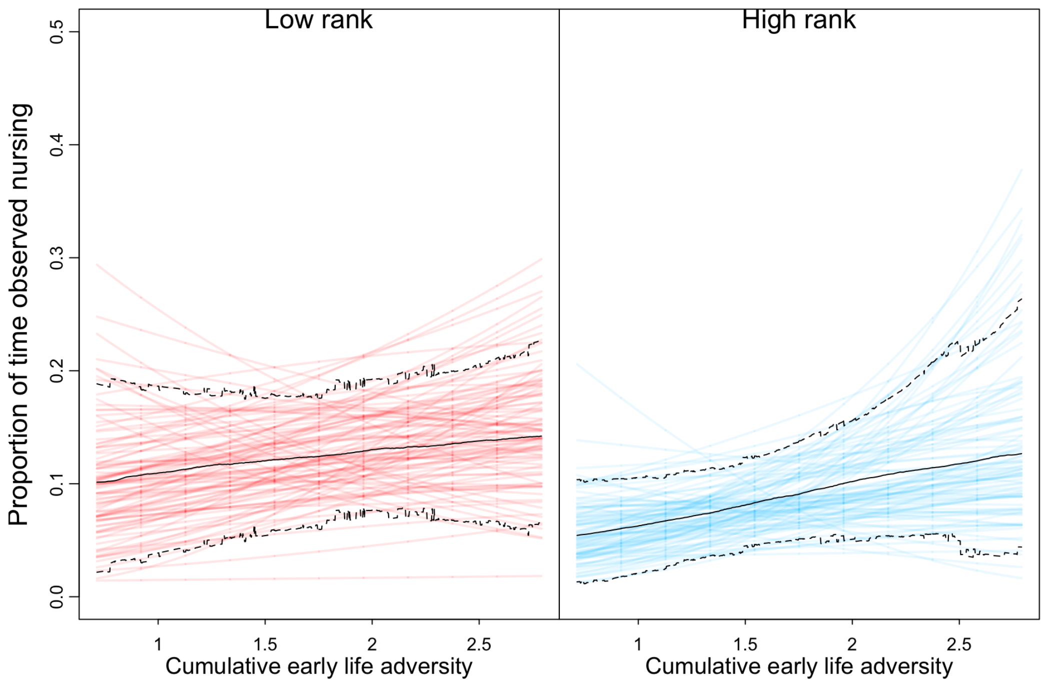


Figure S11. Carrying and early life adversity by maternal dominance rank

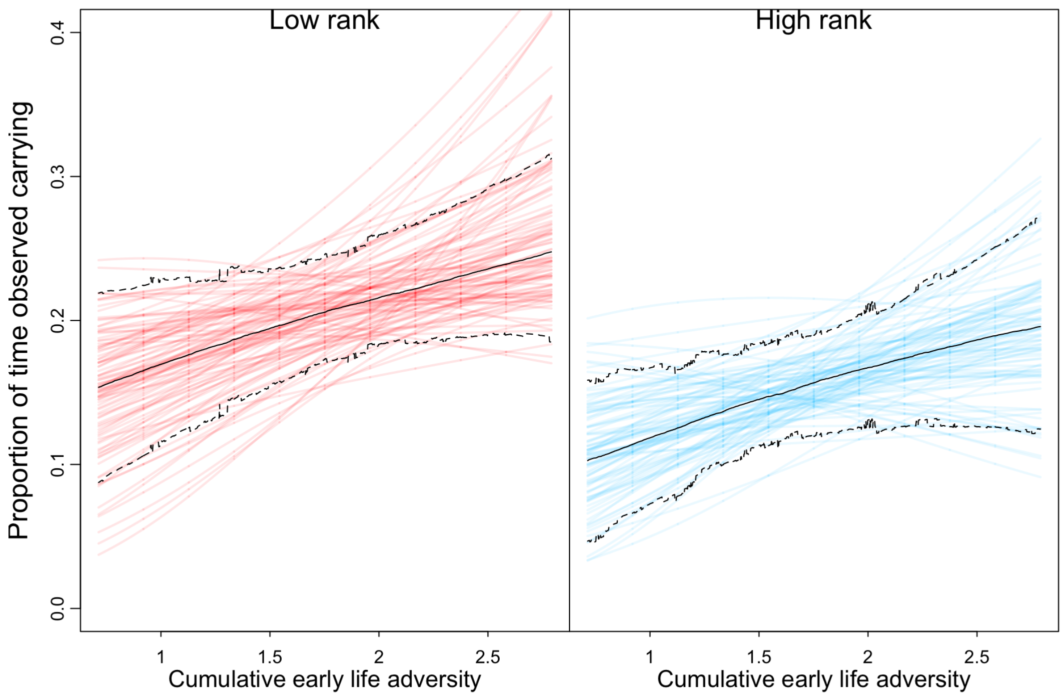

Figure S12. GCMs and early life adversity by maternal dominance rank

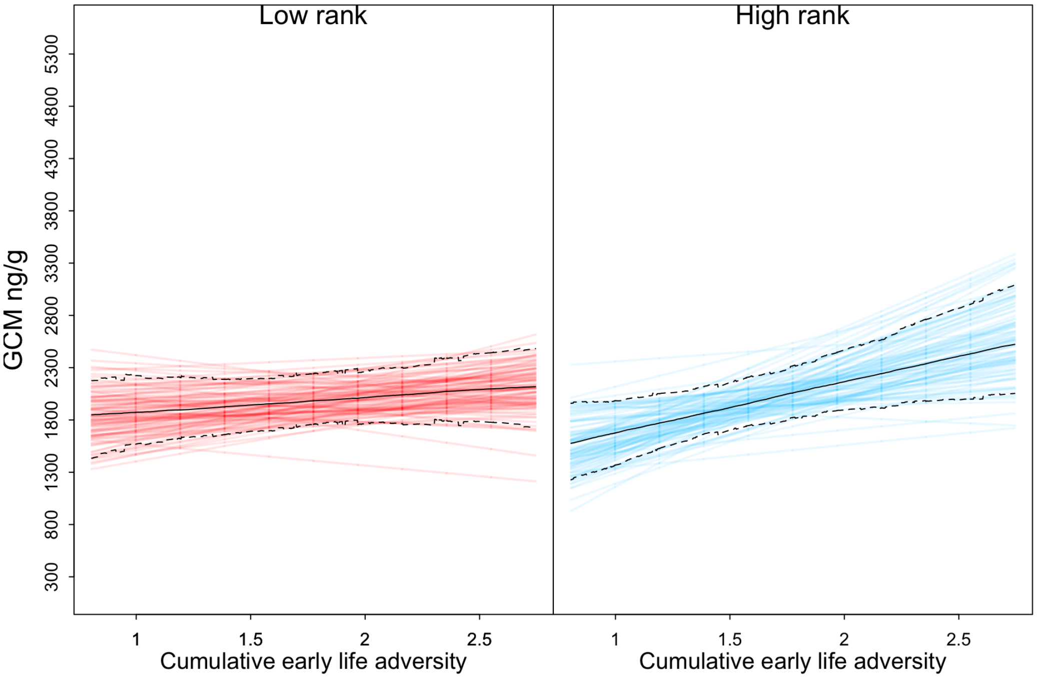


Figure S13. Offspring mortality and early life adversity by maternal dominance rank

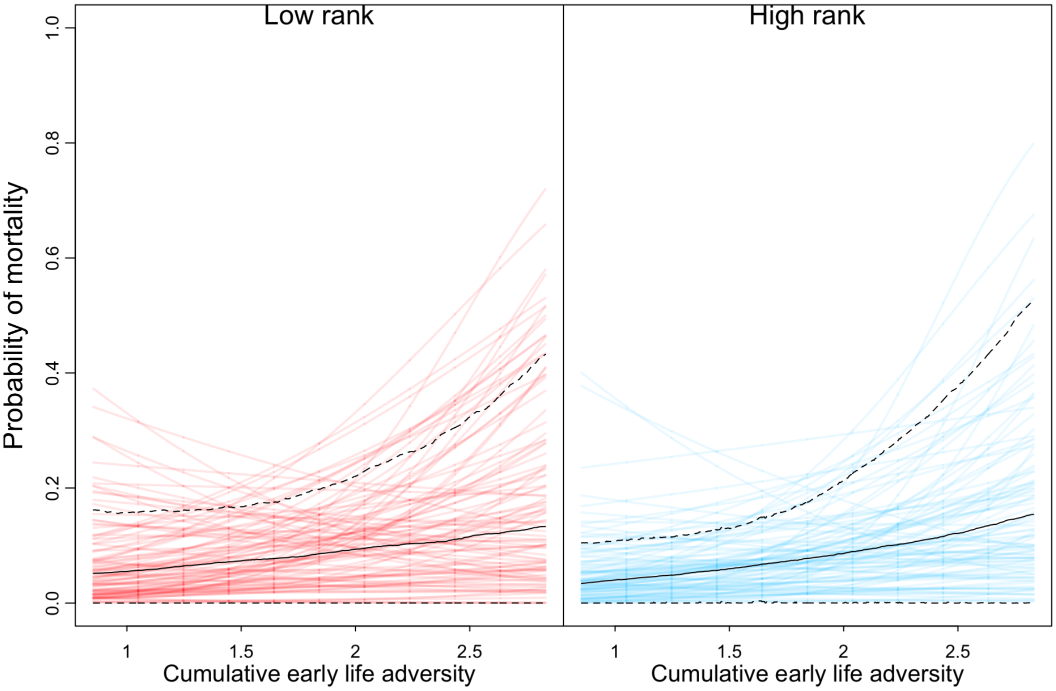
